# Temporal perturbation of STAT1/2 activity reveals dynamic ligand discrimination of type I interferon signaling

**DOI:** 10.1101/2023.03.27.534340

**Authors:** Haowen Yang, Thomas van de Kreeke, Laura C. Van Eyndhoven, Jurjen Tel

## Abstract

Type-I interferon (IFN-I) subtypes signal through the same IFNα receptor (IFNAR), and initiate temporal STAT1/2 activation to orchestrate innate and adaptive immunity. It remains unknown how IFNAR discriminates between subtypes (e.g., IFNα and IFNβ), and how STAT1/2 signaling is affected by time-varying inputs. Here, we utilize our microfluidic system and live-cell imaging to quantify STAT1/2 activation dynamics in a reporter fibroblast model. Population-averaged and single-cell analyses reveal distinct STAT1/2 responses to various IFNα and IFNβ inputs. Upon continuous stimulation, cells show less sensitivity but more sustained responses to IFNα over IFNβ. A short IFNα pulse induces nearly homogeneous STAT1/2 dynamics, in contrast to heterogeneous responses in IFNβ-pulsed cells. Distinct STAT1/2 refractory states emerge upon exposure to repeated IFN-I pulses, while alternating pulse stimulation reveals that IFNβ can revoke STAT1/2 refractoriness caused by IFNα, but not vice versa. These findings highlight the differences between IFNα and IFNβ signaling and how they can elicit distinct temporal cellular behaviors during viral infection.

## INTRODUCTION

Cells constantly encounter multitudes of time-varying environmental cues and stimuli. To process these and act accordingly, individual cells use complex signaling networks to determine their fates and functions (Sears and Nevins, 2002; Hosokawa and Rothenberg, 2021; De Belly et al., 2022). The information of these temporal inputs can be discerned and encoded in temporal patterns of signaling dynamics, such as transcription factor dynamics (Purvis and Lahav, 2013; Sonnen and Aulehla, 2014). Cells also have evolved mechanisms to discriminate myriads of different input molecules to make informed decisions and carry out numerous functions in an orderly fashion. These distinct input signals are usually recognized and transmitted through different receptors, but converge on same signaling pathways to induce specific dynamic responses (Wang et al., 2022). For instance, stimulation with sustained epidermal (EGF)/nerve (NGF) growth factor leads to more transient/sustained ERK activation dynamics (Ryu et al., 2015). NF-κB dynamics can distinguish self (tumor necrosis factor-α, TNF-α) from pathogen (lipopolysaccharide, LPS) signals (Kellogg et al., 2015). Intriguingly, cells also use Notch dynamics to discriminate signaling between the ligands Dll1 and Dll4 through the same receptor Notch1 (Nandagopal et al., 2018).

Type-I interferons (IFN-Is), such as IFNα and IFNβ, play a pivotal role in antiviral and antibacterial immunity (McNab et al., 2015; Van Eyndhoven et al., 2021), antitumor responses (Zitvogel et al., 2015; Zhou et al., 2020), and autoimmune manifestations (Hall and Rosen, 2010; Psarras et al., 2017). To manage such diverse, complex behaviors, cells must be able to discern and encode multitudinous temporal inputs of IFN-I subtypes into distinct signaling dynamics. How cells orchestrate these distinct dynamics is crucial for making informed fate decisions. In vivo, hematopoietic cells are the main source of IFNα, whereas non-hematopoietic cells, such as epithelium, endothelium and fibroblasts, are the main producers of IFNβ (Ivashkiv and Donlin, 2014). In order to establish systemic antiviral immunity, communication between hematopoietic and non-hematopoietic cells is key, while both are susceptible for viral infection. Intriguingly, upon infection, IFNα and IFNβ bind and activate a shared heterodimeric IFNα receptor (IFNAR), which leads to the activation of receptor-associated kinases Janus kinase 1 (JAK1) and tyrosine kinase 2 (Tyk2), regulating the phosphorylation of signal transducer and activator of transcription 1 (STAT1) and STAT2. The phosphorylated STAT proteins homo- or heterodimerize and translocate to the nucleus, promoting the transcription of IFN-stimulated genes (ISGs) to initiate an antiviral state in both infected, and yet uninfected cells ((Ivashkiv and Donlin, 2014; Michalska et al., 2018); Figure 1A).

**Figure 1.**
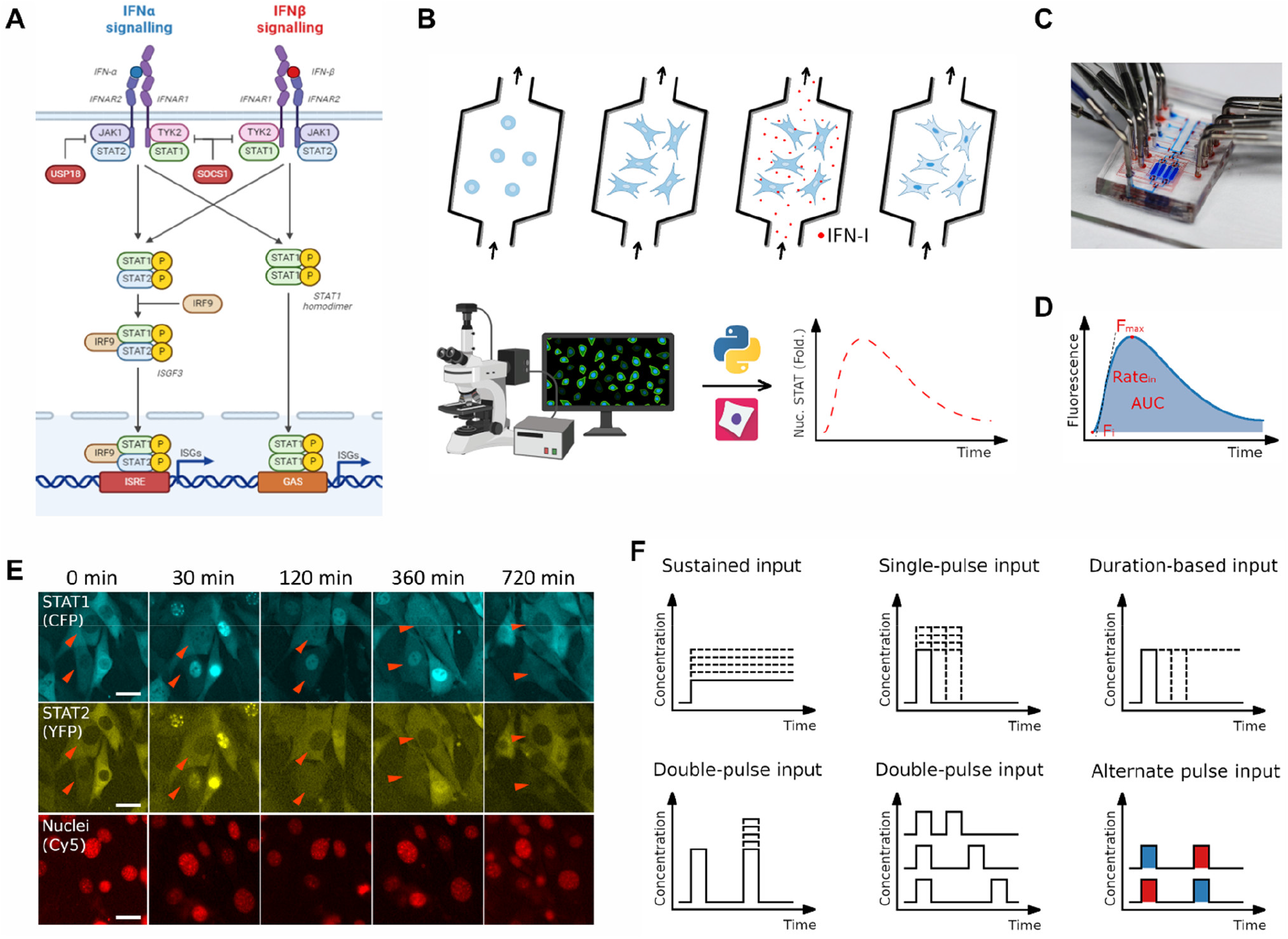
Microfluidic-based approach allows the study of STAT1/2 responses to dynamic signaling inputs. (A) schematic illustrates IFN-I-mediated JAK-STAT1/2 signaling pathway. STAT1 and STAT2 are activated upon IFNα and IFNβ binding IFNAR. (B) The workflow to quantitatively analyze STAT1/2 nuclear translocation activities using flow-based microfluidic approach, live-cell imaging, and data processing. (C) The multilayer microfluidic device integrated with a control layer (red) and a flow layer (blue) that allows cell culture and temporal input delivery. (D) Time course descriptors used to quantify STAT1/2 response dynamics. F_i_ and F_max_ describe the initial and maximal intensity of nuclear STAT fluorescence. Rate_in_ quantifies the maximal rate of STAT nuclear translocation. AUC, the area under the curve of STAT response. (E) Time-lapse images of STAT1-CFP and STAT2-YFP fusion proteins stably expressed in NIH3T3 fibroblasts exposed to a continuous/sustained input of 100 ng/mL IFNβ. Nuclei were stained with BioTracker 650 red nuclear dye. White scale bar, 30 μm. (F) Various temporal input modes of IFN-I used to study STAT1/2 responses. The results obtained under each input condition are shown in Figures 2–7, respectively.

Given that IFNα and IFNβ have considerable structural homology and similarities in function (Thomas et al., 2011; Wittling et al., 2021), we were intrigued to uncover whether cells are able to distinguish between the two subtypes, while IFNAR activation is strictly cell type and context-dependent (van Boxel-Dezaire et al., 2006). Despite that, it has remained largely elusive how cells can nonetheless discriminate between STAT1/2 signaling by different IFN-Is via the common receptor and pathway and elicit distinct downstream responses. To explore these unsolved mysteries, we take advantage of our microfluidic platform to precisely modulate dose and duration of various IFNα/IFNβ inputs, and utilize time-lapse microscopy to track and quantify STAT1/2 activity dynamics in fibroblasts (Figure 1B). We show that IFNα and IFNβ generate distinct activation patterns and heterogeneous dynamics of both STAT1 and STAT2 in single fibroblasts subjected to various IFN-I input types, including sustained, single-pulse and double-pulse inputs with adjustable parameters (strength, duration, identity) (Figure 1F). These results suggest that a ligand-specific identity of IFN-I subtypes is encoded in temporal STAT1/2 activation dynamics, enabling cells to activate distinct STAT1/2 target programs, thereby eliciting specific cellular behaviors. Unraveling this discriminatory capacity of cells will build towards an enhanced understanding on how (non-)hematopoietic cells establish systemic (antiviral) immunity and how it can be modulated for therapeutic purposes.

## RESULTS

### Analyzing dynamic ligand discrimination of IFN-I signaling by imaging STAT1/2 dynamics in individual cells subjected to temporally modulated IFN-I inputs

To determine distinct downstream cell fates upon IFNAR signaling, cells must be able to discriminate between STAT1/2 signaling evoked by temporal profiles of different IFN-Is (e.g., IFNα and IFNβ). To probe the discriminatory capacity of cells, we compared the dynamics of STAT1/2 signaling induced upon various IFNα and IFNβ inputs. To visualize STAT1/2 signaling, we employed NIH3T3 fibroblasts that stably express STAT1-CFP and STAT2-YFP fusion proteins. Upon IFN-I stimulation, the fusion proteins translocate from cytoplasm to the nucleus (Figure 1A), which can be imaged and tracked using time-lapse fluorescent microscopy (Kellogg *et al*., 2015; Yang et al., 2022; Lane et al., 2019).

To examine STAT1/2 responses to various IFN-I input modes, we set up a flow-based, software-programmable microfluidic cell culture system to enable precise modulation of STAT1/2 activity with defined temporal inputs, including continuous, single-pulse, and double-pulse inputs with adjustable amplitudes and durations (Figures 1B, 1C and 1F). Continuous treatment with 100 ng/mL of either IFNα or IFNβ led to nuclear translocation of both STAT1 and STAT2 in the fibroblasts (Figures 1E and S1), confirming the functionality of our platform. Two descriptors, fold change (Lee et al., 2014; Zhang et al., 2017) and nucleus/cytoplasm ratio (Lane *et al*., 2019), are most frequently used in quantifying transcription factor dynamics. Calculation of the coefficient of variation for these two descriptors showed significantly (p<0.05) less variability for fold change (F_max_/F_i_) (Figures 1D and S2A), which was selected for the quantification of STAT1/2 dynamics. Quantitative evaluation of peak amplitudes suggested a strong correlation of STAT1 activity with STAT2 dynamics (Figure S2B).

Together, the combination of our microfluidic system and time-lapse imaging allows quantitative analysis of STAT1/2 dynamics in individual cells subjected to various temporal IFN-I inputs.

### Continuous stimulation with IFNα and IFNβ induce distinct, heterogeneous STAT1/2 activation dynamics

To quantify the STAT1/2 signaling dynamics induced upon continuous IFNα or IFNβ inputs, we stimulated cells with each IFN-I subtype of different concentrations ranging from 0.1 to 100 ng/mL. The population-average results showed that high-dosage IFNα triggered a single STAT1 activation peak that gradually dropped to baseline after 8 h, while it evoked a STAT2 activation peak with a sharp decline after the first 2 h and a slow return to baseline afterwards. Similar STAT1/2 dynamics profiles were observed upon stimulation with high-dosage IFNβ (Figure 2A). However, low-dosage IFNβ was still able to induce stronger STAT1/2 responses while a same dose of IFNα was not (Figure 2A), as demonstrated by peak amplitudes and rate of nuclear change (rate_in_) of STAT1/2 (Figures S3B and S3C). A similar trend was observed in the fraction of responding cells (STAT1/2 activation rate) for both IFNα and IFNβ, where low-dosage IFNβ activated a significantly greater proportion of cells compared to the same dose of IFNα (Figure 2B). Using the activation rates, we calculated a dose-response curve, resulting in a half maximal effective concentration (EC_50_) of 3.26 and 0.0573 ng/ml for IFNα and IFNβ, respectively. This indicated that IFNβ activated 50% of cells with a dose of 57 times lower than IFNα, which means cells were more sensitive to IFNβ over IFNα. The median value of STAT1/2 peak response time showed a monotonic decrease over IFNβ dose, while increased in the range of low-dosage IFNα from 0.1 to 1 ng/mL followed by a decline as the dose kept rising (Figure S3D).

**Figure 2.**
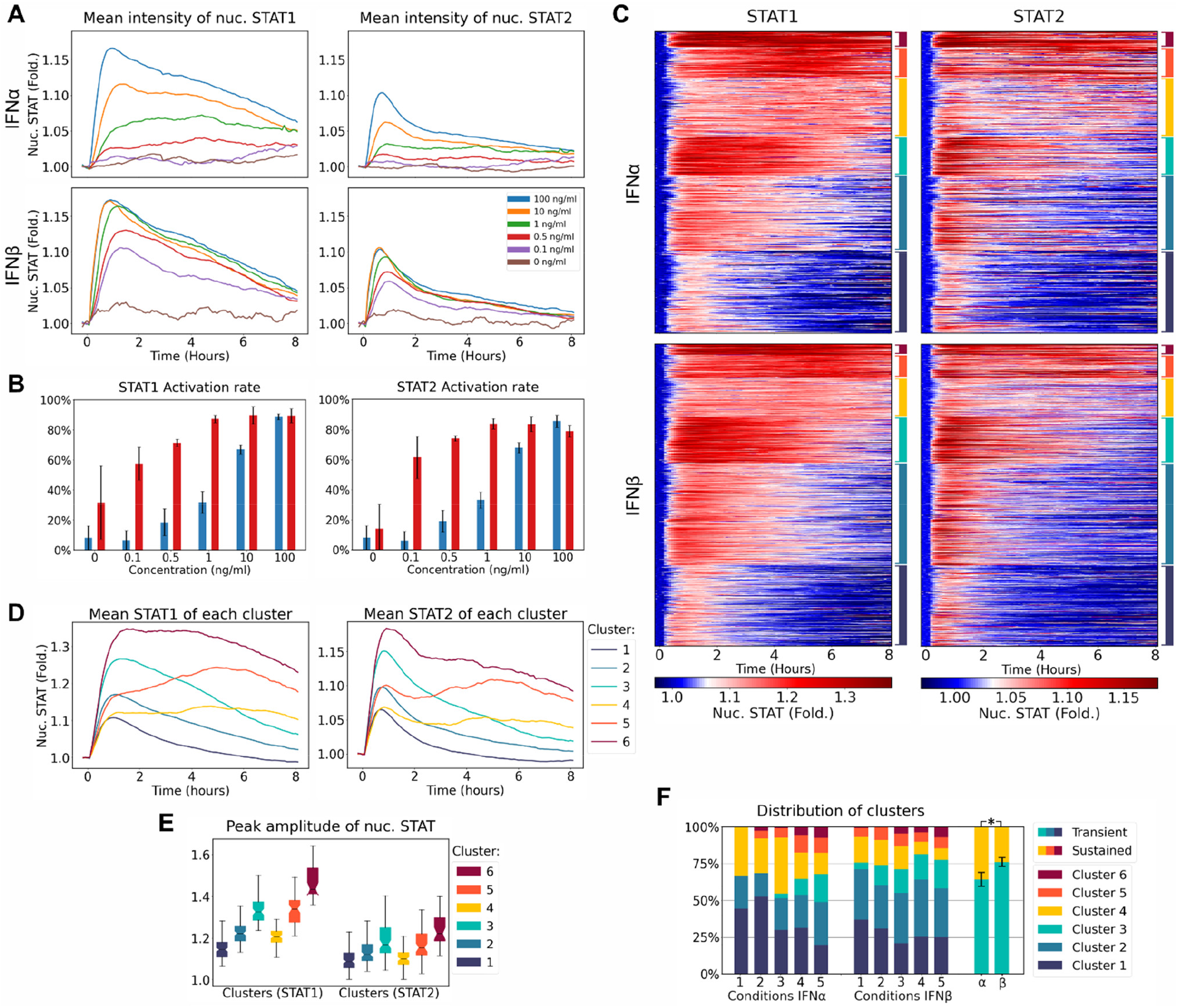
STAT1/2 responses to continuous IFN-I inputs. (A) Population-average STAT1/2 nuclear translocation dynamics upon continuous stimulation with different concentrations of IFNα/IFNβ ranging from 0.1–100 ng/mL (0 ng/mL as a blank control). (B) Fraction of responding cells after continuous stimulation under the same conditions in (A). (C) Clustered single-cell STAT1/2 activity trajectories under the conditions in (A) were displayed in heatmaps. The negative cells were excluded. (D) Average STAT1/2 activity across 6 clusters identified in (C). (E) Peak amplitudes of single-cell STAT1/2 trajectories across 6 clusters from (C). (F) Distribution of STAT1/2 activity across 6 clusters from (C) and (D) in response to different IFN-I dosages.

The single-cell trajectory datasets showed heterogeneous STAT1/2 dynamics for both IFN-I subtypes (Figure S3A). To quantify the heterogeneity, we pooled all doses of IFNα- and IFNβ-induced STAT1/2 activation trajectories and used a self-organizing maps (SOM) clustering algorithm (Tamayo et al., 1999; Ong et al., 2021) to extract 6 temporal activation patterns (Figure 2C). At the single-cell level, we observed a mix of transient and sustained STAT1/2 dynamics present upon both IFNα- and IFNβ-induced signaling, showing heterogeneous STAT1/2 dynamics with a similar distribution of each cluster between IFNα and IFNβ (Figures 2C–2E). To investigate which cellular response pattern originated from which input dose, we calculated the incidence of each cluster in response to different IFN-I doses (Figures 2E and 2F). Intriguingly, we found that both IFNα and IFNβ favored transient responses, while IFNα induced a significantly larger fraction of sustained STAT1/2 activation pattern than IFNβ under each condition (Figures 2D and 2F). In contrast to STAT1 trajectory clustering, STAT2 sustained clusters showed a sharp peak followed by a plateau (Figure 2D), potentially indicating bistability. Another interesting observation was that the medium IFN-I dose (1 ng/mL) shifted the distribution of STAT1/2 activity trajectories toward more sustained profiles when compared to both the high and low IFN-I doses (Figure 2F). Importantly, such shifts were inconspicuous in population-average dynamics (Figures 2A and S3A).

Together, these results demonstrate that cells upon constant IFN-I exposure showed distinct STAT1/2 signaling by IFNα compared to IFNβ, where IFNα showed higher EC_50_ value on STAT1/2 activation but induced a larger fraction of sustained STAT1/2 dynamics in comparison with IFNβ.

### Single-pulse IFNα and IFNβ can be discriminated by population-averaged peak response time and single-cell heterogeneity on STAT1/2 dynamics

Since cells encode input information from both the amplitude and duration of inputs (Kellogg *et al*., 2015), we applied single pulses of IFNα or IFNβ with varying concentrations and/or durations to the fibroblasts. Intriguingly, the averaged responses showed that changing IFN-I (IFNα or IFNβ) concentration under constant duration maintained timely STAT1/2 peak responses (Figure 3A). Notably, in contrast to IFNβ, increasing IFNα pulse duration led to greater delay in STAT1/2 peak responses (Figure 3C). The peak amplitude from different conditions followed the same pattern as shown in the averaged STAT1/2 dynamics (Figures 3A, S4A and S4B). The longer IFN-I pulse resulted in greater STAT1/2 response duration (Figure S4C), and larger area under the curve (AUC) (Figure S4D), suggesting that input duration determines response duration. Similar to continuous stimulation, a single pulse of IFNβ (from 50 to 100 ng/mL) maintained nearly constant STAT1/2 activation rates due to its higher EC_50_ value, while single IFNα pulses resulted in an increasing number of active cells over input concentration (Figure 3B).

**Figure 3.**
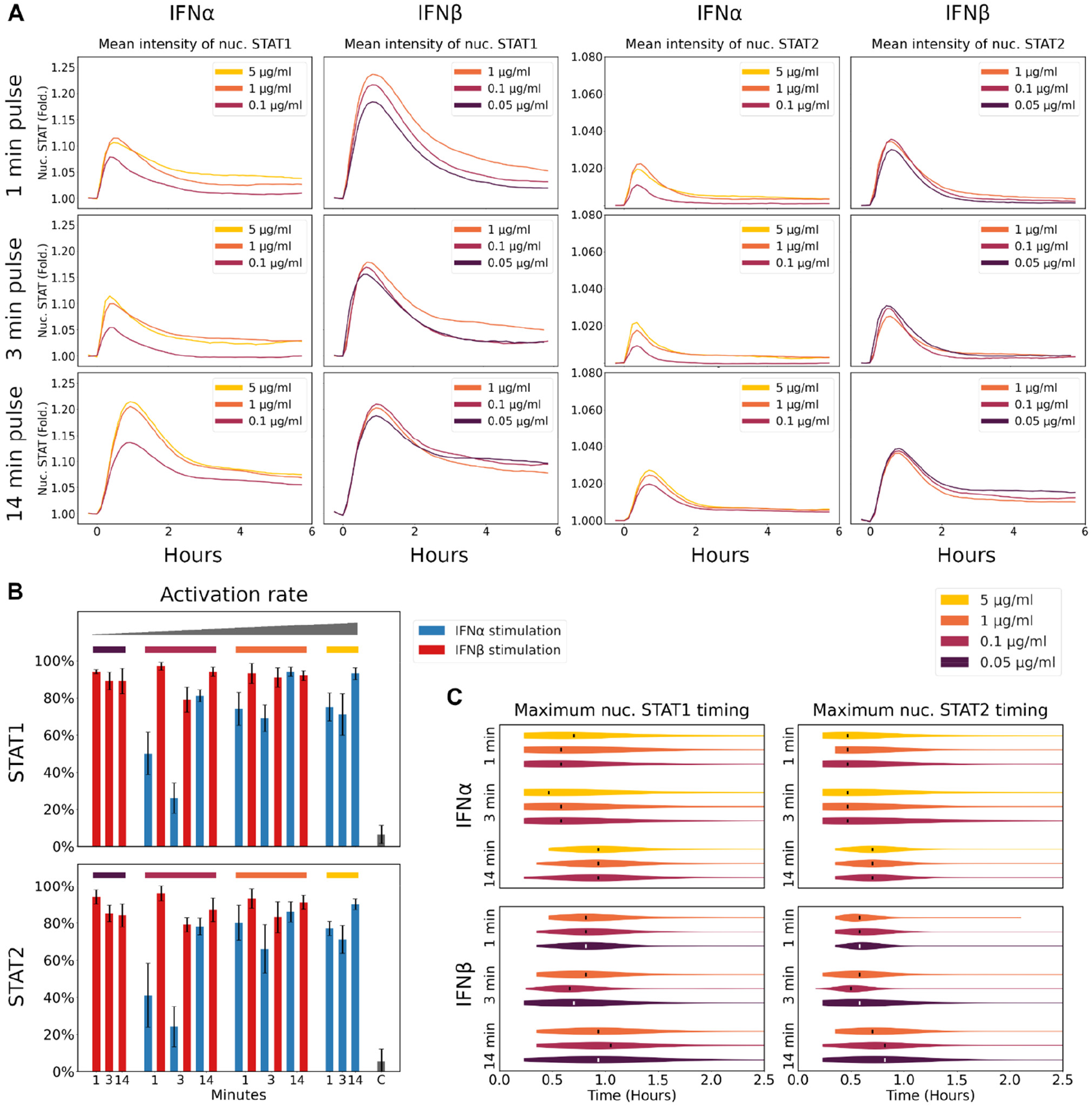
STAT1/2 responses to single-pulse IFN-I inputs. (A) Population-average STAT1/2 activation dynamics upon single-pulse IFNα/IFNβ stimulation at different concentrations (0.1, 1 and 5 μg/mL) or durations (1, 3, and 14 min). (B) Fraction of responding cells after single-pulse stimulation under the conditions in (A). (C) Peak response timing of single-cell STAT1/2 activity trajectories under the conditions in (A).

To further compare single-cell heterogeneity on STAT1/2 dynamics by single-pulse IFN-I subtypes, we pooled all STAT1/2 activity trajectories by 100 ng/mL IFNα or IFNβ with all input durations (1, 3, 14 min, and continuous) together, and extracted 6 clusters using the SOM algorithm (Figures 4A and 4B), illustrating that sustained activation patterns became more apparent over input duration (Figure 4A). These results demonstrated that IFN-I input duration controls cellular heterogeneity on STAT1/2 responses. Noticeably, we found that upon a short IFNα pulse stimulation cells showed nearly homogeneous STAT1/2 responses, while a single-pulse IFNβ induced greater heterogeneity on STAT1/2 dynamics (Figure 4C). Moreover, a medium IFNβ pulse duration (14 min) led to a much greater heterogeneity on STAT2 when compared to IFNα application (Figures S5A–S5C). On average, 22% of cells sustainedly responded to a 14 min IFNβ pulse while only 5% of active cells showed long-lasting STAT2 activation when exposed to a same pulse duration of IFNα (Figures S5B and S5C). In addition, the bistable STAT dynamics were observed neither in short or in medium pulse IFN-I-treated cells, possibly indicating that bistable behavior of cells were triggered only upon long-duration input (Figures 4B and 4C, S5B and S5C).

**Figure 4.**
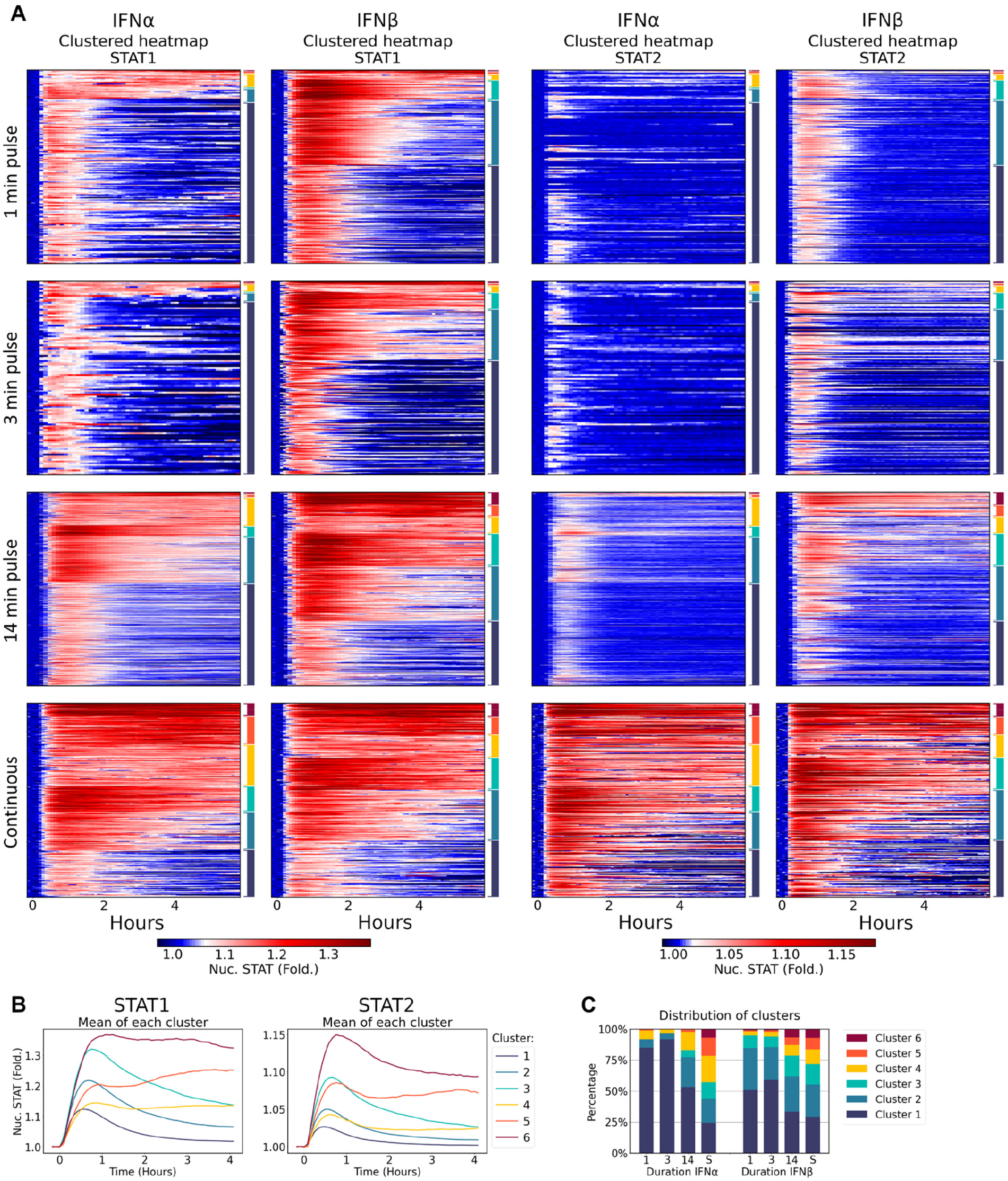
STAT1/2 responses to duration-based IFN-I inputs. (A) Clustered single-cell STAT1/2 activity trajectories upon 100 ng/mL IFNα/IFNβ stimulation at different durations (1, 3, 14 min and continuous) were displayed in heatmaps. The negative cells were excluded. (B) Average STAT1/2 activity across 6 clusters identified in (A). (C) Distribution of STAT1/2 activity across 6 clusters from (A) and (B) in response to different IFN-I durations.

Altogether, cells displayed distinct features of STAT1/2 signaling between IFNα and IFNβ upon single-pulse stimulation, such as averaged STAT1/2 peak response time and STAT1/2 activity trajectory at single-cell level.

### Repeated pulse stimulation reveals that IFNα and IFNβ evoke distinct STAT1/2 peak amplitude saturation, heterogeneity and refractory states

Prestimulation with high-dosage IFNα was reported to desensitize the STAT1/2 pathway (Kok et al., 2020). We suspected that IFNβ probably has a similar effect, which we aimed to unravel. Besides, we were also interested in whether repeated pulse stimulation with IFN-I subtypes trigger distinct STAT1/2 activation dynamics. To address these questions, we applied repeated 3 min pulses of 100 ng/mL IFNα or 10 ng/ml IFNβ that initiated double peak STAT1/2 responses (Figures 5A and 5B). Restimulating cells with the same IFN-I concentration resulted in a slight decrease in averaged STAT1/2 nuclear translocation. Increasing the strength of the second IFN-I pulse triggered a stronger STAT1/2 reactivation. The second STAT1/2 peak amplitude saturated above a concentration 10 and 5 times the first pulse of IFNα and IFNβ, respectively (Figure 5A). These results indicated the desensitization of STAT1/2 pathway to both IFNα and IFNβ. Furthermore, the fraction of cells responding to a second input pulse dropped by around 10% in the second STAT1/2 responses (Figure 5C). Cells showed a higher EC_50_ dose response to a second pulse of IFNα compared to IFNβ after the first pulse stimulation with the same IFN-I (Figure 5C), as similarly found in the responses under continuous stimulation.

**Figure 5.**
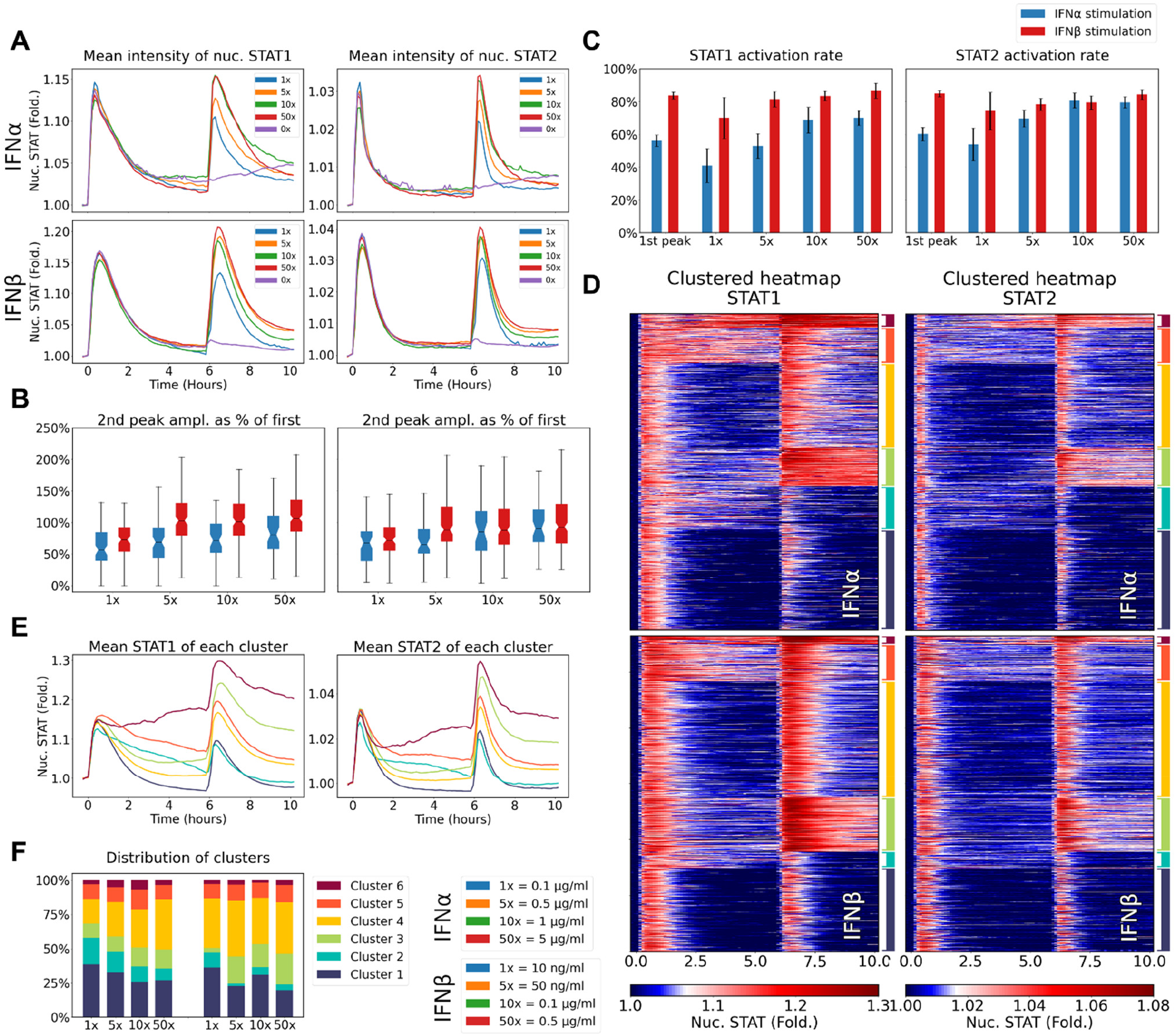
STAT1/2 responses to repeated IFN-I pulses with different intensities at the second pulse. (A) Population-average STAT1/2 activity dynamics upon exposure to repeated IFNα or IFNβ pulses, with a first pulse (3 min) at a concentration of 0.1 or 0.01 μg/mL, and a second pulse (3 min) ranging from 0.1–5 or 0.01–0.5 μg/mL. (B) Fraction of responding cells after the first and second pulse stimulation under the conditions in (A). (C) Clustered single-cell STAT1/2 activity trajectories under the conditions in (A) were displayed in heatmaps. The negative cells were excluded. (D) Peak amplitudes of single-cell STAT1/2 trajectories across 6 clusters from (C). (E) Average STAT1/2 activity across 6 clusters identified in (C). (F) Distribution of STAT1/2 activity across 6 clusters from (C) and (E) in response to different IFN-I concentrations of second pulse.

To quantify the magnitude of the peak STAT1/2 responses, we calculated the second peak amplitude as a percentage of the first one (Figure 5C). Stimulation with a repeated pulse of IFNα or IFNβ showed an attenuated second peak as 62% or 75% of the first one on STAT1 responses, with a similar observation on STAT2 responses. When re-pulsed with IFNβ with a concentration of 5, 10 or 50 times the first pulse, the averaged second STAT1/2 peak amplitude increased to 108% of the first peak with no significant difference between these 3 conditions, indicating the saturation of the second STAT1/2 peak amplitude. In contrast, the amplitude of the second peak remained below 100% of the first one on average, indicating that a second pulse of IFNα was not able to reactivate cells as potent as the first-pulse. In addition, we measured the area under the curve (AUC) of STAT1/2 dynamics in individual cells for each condition. AUC2 (AUC of the second response) was normalized by AUC1 (AUC of the first response) for the same cell (AUC2/AUC1), and plotted against AUC1, showing a clear separation between conditions in proportion with the concentration of second IFN-I pulse (Figure S6A). To compare cells with similar responsiveness to IFN-I, data were binned by AUC1 and we plotted AUC2 against input pulse concentration. Logarithmic curves were fitted to these datapoints, illustrating that STAT1/2 peak amplitudes became saturated over the concentration of 5 μg/mL for IFNα, which was 50 times those triggered by IFNβ (0.1 μg/mL) (Figure S6B). This indicated that STAT1/2 peak amplitudes saturated faster with increasing IFNβ concentrations at the second pulse compared to those upon repeated IFNα stimulation.

We further explored single-cell heterogeneity on STAT1/2 dynamics upon exposure to double-pulse IFN-Is (Figures 5D–5F). We identified 6 clusters, with 4 clusters resembling in-phase STAT1/2 activity trajectory (clusters 1, 3, 4 and 5) and 2 resembling out-of-phase STAT1/2 dynamics (clusters 2 and 6) (Figures 5D and 5E). We found 3 in-phase clusters (1, 4 and 5) stably presented in both IFNα- and IFNβ-restimulated cells under each condition, as well as 1 out-of-phase response (cluster 6) despite its low incidence. However, in marked contrast to IFNα restimulation, an out-of-phase (cluster 2) trajectory was favored upon the second pulse of IFNβ input with a same strength as the first pulse, while an in-phase (cluster 3) became dominant when the strength of the second pulse was largely increased (Figures 5E and 5F). This interesting observation indicated that a small subset of cells exhibited differential responses to double-pulse IFNα and IFNβ exposure, and implies the ability of IFNβ to adjust the distribution of in-phase and out-of-phase responses.

When cells encounter short pulse intervals, some pathways (such as ERK and NF-κB) barely respond to a second pulse input (such as EGF/NGF and TNFα) and thus display refractory states (Ryu *et al*., 2015; Adamson et al., 2016). We asked whether STAT1/2 activity shows distinct refractory states upon treatment with double IFN-I pulses. To explore the timescale at which STAT1/2 reactivation occurs, we applied double-pulse IFNα (100 ng/mL) or IFNβ (20 ng/mL) at time intervals ranging from 2 to 8 h. The average data showed that cells were able to respond to a second pulse of IFNα or IFNβ upon all the time interval conditions (Figure 6A). Strikingly, while most of the cells responded to the first IFN-I pulses in both situations, cells largely failed to respond to the second pulse at each pulse interval condition, showing a refractory state in the STAT1/2 activity. For example, at a 2 h pulse interval, only 11% and 2% of cells responded to a second IFNα and IFNβ pulse, respectively (Figure 6B). Surprisingly, upon repeated IFNα stimulation at a 4 h interval, the proportion of responding cells increased to 26% but decreased sharply again at longer intervals; while the fraction of active cells monotonically increased over IFNβ pulse interval and maintained stable at 6 h (23%) and longer (25%) intervals (Figure 6B). The non-monotonic trend of IFNα-induced reactivation rate versus pulse intervals might be attributed a delayed negative feedback that only inhibited the response to IFNα but not IFNβ, such as ubiquitin specific peptidase 18 (USP-18)-based negative regulation (Wilmes et al., 2015; Arimoto et al., 2018; Vizan et al., 2013). These data further indicated that the measurement of population-averages masked variability in STAT response at the single-cell level. Furthermore, the amplitude of the second STAT1/2 response became lower but modestly increased over pulse interval (Figure 6C), indicating a refractory state of STAT1/2 activity and a transient desensitization effect on IFN-I restimulation.

**Figure 6.**
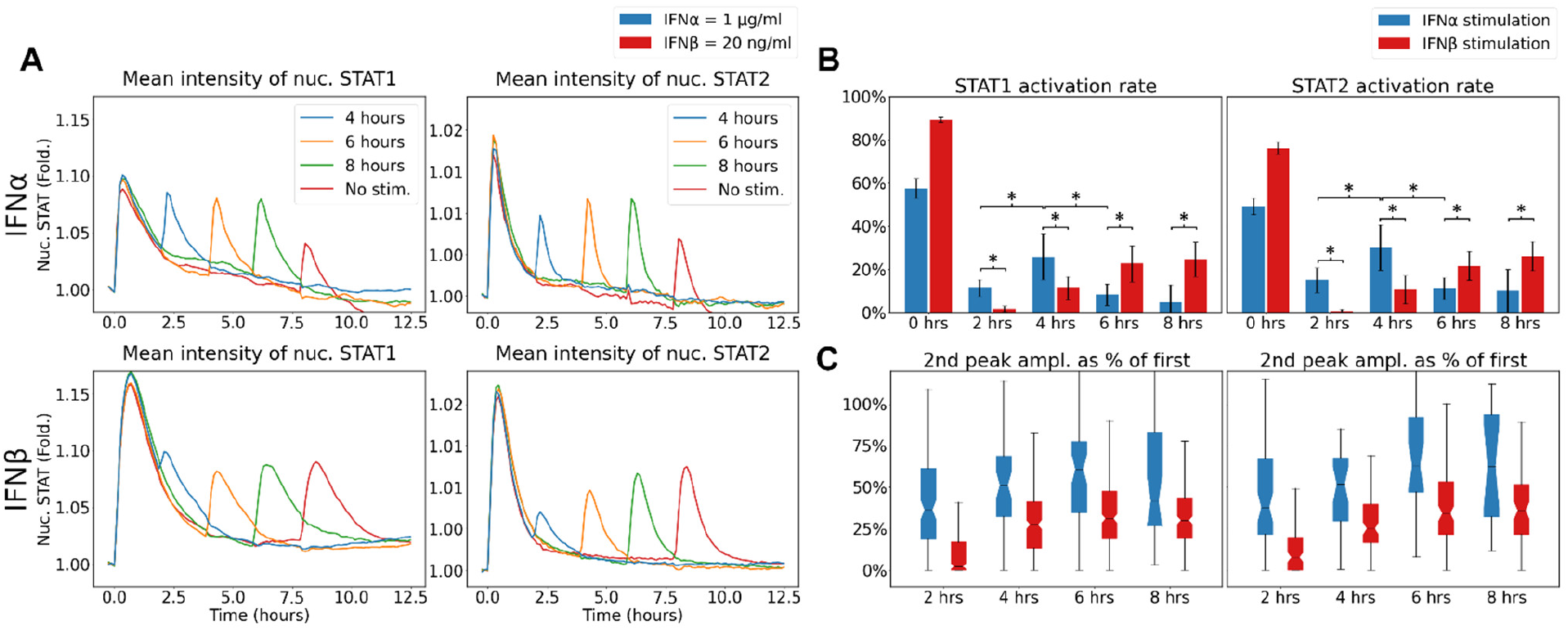
STAT1/2 responses to repeated IFN-I pulses with different time intervals. (A) Population-average STAT1/2 activity dynamics upon exposure to repeated pulses (1 or 0.02 μg/mL IFNα or IFNβ) with 2, 4, 6 and 8 h intervals separately. (B) Fraction of responding cells after the first and second pulse stimulation under the conditions in (A). (C) The second peak amplitude of single-cell STAT1/2 activity trajectories expressed as a fraction of first peak amplitude under the conditions in (A).

Taken together, upon double-pulse stimulation, cells exhibited distinct STAT1/2 dynamics between IFNα and IFNβ, including saturation of the second peak amplitude, single-cell heterogeneity, and refractory state.

### Alternating pulse stimulation reveals that cells refractory to IFNα respond to IFNβ, but not vice versa

TNFα-caused refractory states in the NF-κB system can be revoked by restimulating with interleukin 1β (IL-1β), thus reactivating the NF-κB response (Adamson *et al*., 2016). Such a mechanism may allow cells to use refractory states to discriminate between different cytokines. We were curious whether alternating stimulation with different IFN-Is can revoke STAT1/2 refractory states caused by either IFN-I subtype. We compared responses to alternating pulses of IFNα and IFNβ (αβ) to stimulation with alternate IFNβ and IFNα (βα) pulses at 2 and 6 h intervals separately. First, we applied a 3 min pulse at a same concentration of 50 ng/mL for both conditions. As expected, the peak amplitude of the second STAT1/2 responses became stronger when treated with αβ pulses at both 2 and 6 h intervals, in contrast to stimulation with βα pulses which greatly attenuated the second response (Figure 7A). Also, about 90 or 68% of cells responded when IFNα pulsed-cells were stimulated with an IFNβ pulse after 2 or 6 h, while less than 10% of cells showed STAT1/2 reactivation upon βα pulses stimulation (Figure 7B). In cells subjected to αβ pulses, the average amplitude of the second STAT1/2 nuclear translocation was significantly higher and equated to 120% of the first-peak amplitude. However, upon treatment with βα pulses, the nuclear STAT1/2 amplitude dramatically diminished to 10% of the first-peak amplitude at both pulse interval conditions (Figure 7C). We further doubled the IFNα concentration (100 ng/mL) and maintained the same concentration (50 ng/mL) for IFNβ. We found that the effect of IFNα on the second STAT1/2 response was significantly enhanced in all conditions (Figures 7D–7F). For example, when cells were pulsed by 100 ng/mL IFNα, a smaller fraction of cells responded to an IFNβ pulse after 2 or 6 h in contrast to stimulation with αβ pulses at the original concentration. However, the number of cells pulsed by IFNβ responded to a 100 ng/mL IFNα pulse almost doubled compared to responses to the original βα pulses (Figure 7E). A 100 ng/mL IFNα as either the first or second pulse evoked a reduction in the peak amplitude of the second STAT1/2 response upon either alternate input condition. Nevertheless, upon a second pulse stimulation with 100 ng/mL IFNα, the decrease of STAT1/2 peak amplitude between the first and second responses became smaller compared to treatment with the original βα pulses (Figure 7F).

**Figure 7.**
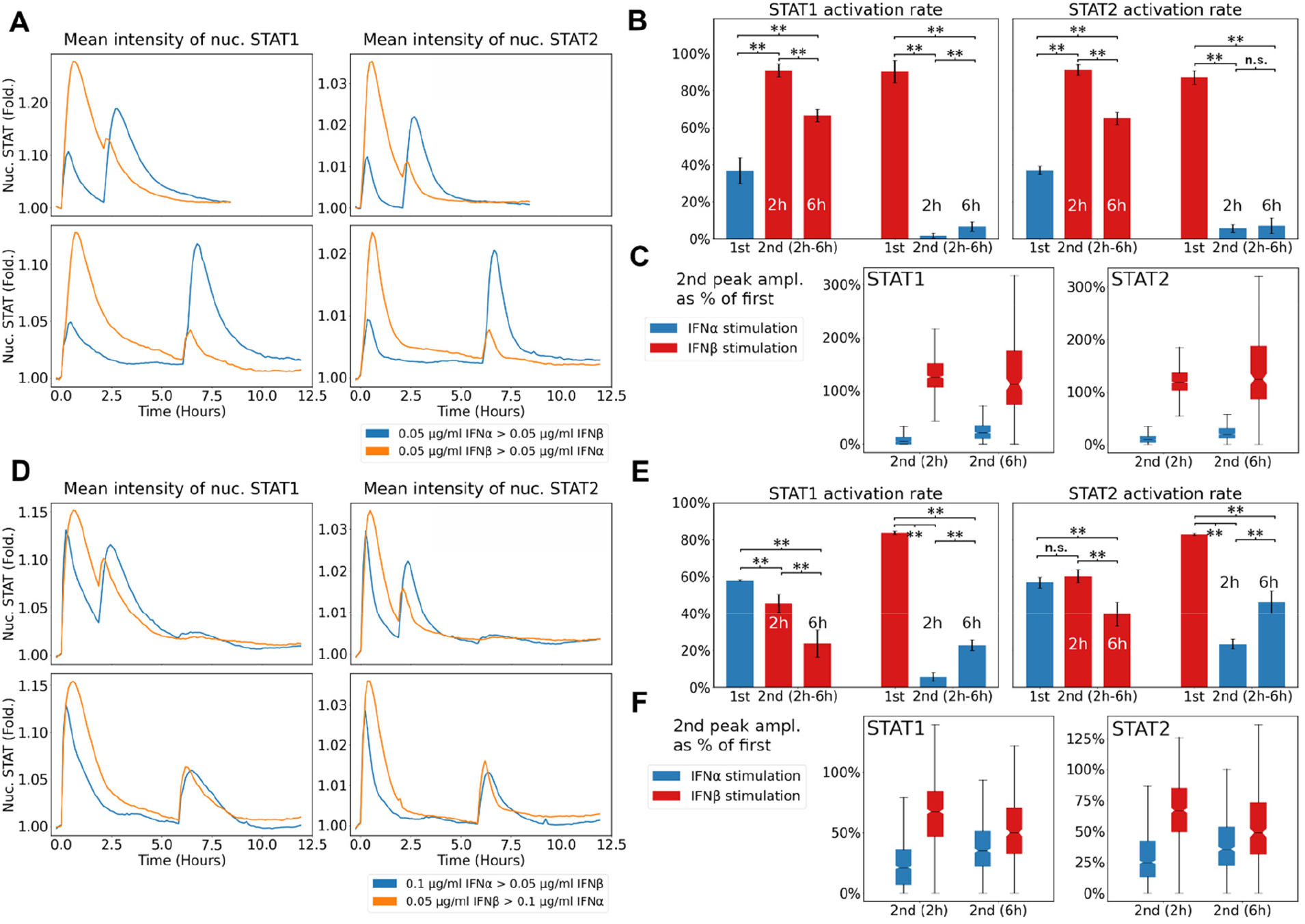
STAT1/2 responses to alternate IFNα and IFNβ pulses. (A) Population-average STAT1/2 activity dynamics upon exposure to alternating pulses at equal intensity, with a 2 or 6 h interval. Blue: STAT1/2 responses to a first pulse with 3min, 0.05 μg/mL IFNα and a second pulse with 3 min, 0.05 μg/mL IFNβ. Orange: STAT1/2 responses to a first pulse with 3 min, 0.05 μg/mL IFNβ and a second pulse with 3 min, 0.05 μg/mL IFNα. (B) Fraction of responding cells after the first and second pulse stimulation under the conditions in (A). (C) The second peak amplitude of single-cell STAT1/2 activity trajectories expressed as a fraction of first peak amplitude under the conditions in (A). (D) Population-average STAT1/2 activity dynamics upon exposure to alternating pulses at different intensity, with a 2 or 6 h interval. Blue: STAT1/2 responses to a first pulse with 3min, 0.1 μg/mL IFNα and a second pulse with 3 min, 0.05 μg/mL IFNβ. Orange: STAT1/2 responses to a first pulse with 3 min, 0.05 μg/mL IFNβ and a second pulse with 3 min, 0.1 μg/mL IFNα. (E) Fraction of responding cells after the first and second pulse stimulation under the conditions in (D). (F) The second peak amplitude of single-cell STAT1/2 activity trajectories expressed as a fraction of first peak amplitude under the conditions in (D).

Altogether, these results revealed that IFNβ stimulation can revoke the STAT1/2 refractory state caused by IFNα, but not vice versa.

## DISCUSSION

It is well established that IFN-I subtypes (e.g., IFNα and IFNβ) signal through the same receptor IFNAR and trigger distinct downstream responses. However, it has remained largely unknown how cells discriminate between these IFN-I subtypes via the same receptor. One possible strategy is encoding different IFN-Is into distinct transcription factor dynamics (Nandagopal *et al*., 2018; Thomas *et al*., 2011). In this study, we investigated the role of STAT1/2 activity dynamics on ligand discrimination of IFN-I signaling to include the processing of diverse temporal input types (Figure 1E). Measuring activation dynamics of STAT1 and STAT2 in individual cells revealed the capacity of JAK-STAT signaling system to discriminate between IFN-I subtypes under various input modes. These findings add support to the growing body of literature on how cells use transcription factor activation dynamics to encode distinct ligand information about different temporal inputs.

Important questions arise why IFN-I subtypes are necessary and how they can be discriminated and induce different cell fates when Figurehting pathogen invasion. One possible reason could be the different sources of IFN-I subtypes. IFNα is mostly produced by hematopoietic cells (Ivashkiv and Donlin, 2014; Ali et al., 2019), where plasmacytoid dendritic cells (pDCs) are the predominant producer (Bencze et al., 2021), whereas IFNβ is predominantly produced by structural cells, such as epithelium, endothelium and fibroblasts (Krausgruber et al., 2020). The expression patterns of immune genes modulated the extensive interactions between structural cells and hematopoietic immune cells in mice undergoing systemic viral infection (Krausgruber *et al*., 2020). Although most viruses infect structural cells (such as fibroblasts) instead of pDCs, pDCs can produce IFNα without being infected themselves (Bencze *et al*., 2021). In addition, IFNβ priming facilitates massive IFNα production by pDCs once infection occurs (Wimmers et al., 2018). Fibroblasts can also be primed with IFNα that is constitutively produced by pDCs or other myeloid cells. After being infected, these primed fibroblasts will produce large amounts of IFNβ (Krausgruber *et al*., 2020). In addition, fibroblasts or other structural cells may respond to high levels of alternating IFNα and IFNβ waves in different situations dependent on the process of infection. For example, infected fibroblasts produce a huge amount of IFNβ, and further activates pDCs upon interferogenic synapse formation (Assil et al., 2019), consequently leading to massive IFNα production by pDCs. Being infected, localized fibroblasts activate pDCs via interferogenic synapses to produce large amounts of IFNα that will further stimulate distal fibroblasts. As virus spreads, the distal fibroblasts become infected and subsequently result in a big wave of IFNα. We hypothesize that distinct cell fates (states of STAT1/2 activation dynamics) are determined by differential encoding of these IFN-I subtypes at various temporal conditions.

Analyzing STAT1/2 responses at single-cell level not only allows us to observe different activation states but also signaling dynamics in individual cells. We demonstrated that STAT1/2 signaling response to IFNα and IFNβ are distinct under each input mode. At sustained input mode, both IFNα and IFNβ led to switch-like (also known as digital) activation patterns (Kellogg *et al*., 2015), in spite of a 57 times lower EC_50_ value in STAT activation rate for IFNβ compared to IFNα. At high input doses, the resulting STAT1/2 peak amplitudes became saturated. This overrides the capacity of fibroblasts to discriminate between sustained IFNα and IFNβ, leading to remarkably similar STAT1/2 signaling profiles in the cells, both presenting a mix of sustained and transient responses. This feature possibly arises from the inherent variability in expression levels of signaling proteins in JAK-STAT pathway, and different strengths of positive and negative feedback regulation within different cell individuals (Ivashkiv and Donlin, 2014; Michalska *et al*., 2018; Porritt and Hertzog, 2015). However, at low input doses, the distinct STAT1/2 responses by IFNα and IFNβ became apparent, suggesting that fibroblasts might be more capable of discriminating self IFN-I signal (IFNβ) from non-self input (IFNα) at low dosage level under long-term IFN-I exposure. This phenomenon possibly implies that signal discrimination based on the different potency might be an easy way when discerning highly similar signals (such as cytokine subtypes).

When encountering short pulse input, cells exhibited less potent responses to IFNα than to IFNβ, irrelevant to the input concentration. The longer pulse led to the saturation of STAT1/2 peak amplitudes by IFNβ but was still able to enhance STAT1/2 responses by IFNα. At 14 min pulse, the high input concentration induced an equal STAT1 peak amplitude by IFNα and IFNβ, but a stronger STAT2 response upon IFNβ treatment, which may result from the differential upregulation of STAT2 and IRF9 (Michalska *et al*., 2018; Majoros et al., 2017). Short IFNα pulse maintained nearly homogeneous STAT1/2 responses, in contrast to greater heterogeneity on STAT1/2 dynamics in IFNβ-pulsed cells. These findings emphasize the importance of using low input durations to reveal how individual cells process different IFN-I information.

Repeated pulse stimulation with gradient input concentration led to a slight increase and saturation of STAT1/2 peak amplitudes, even when the strength of second pulse became 50 times that of the original pulse. This is possibly due to the limited levels of cytoplasmic STAT1 and STAT2, resulting in an insufficient reservoir of STAT proteins to afford stronger activation. Clustering of single-cell trajectories showed similar distribution of each STAT1/2 cluster present among all conditions in repeated IFNα stimulation, while when increasing the strength of second-pulse IFNβ, an out-of-phase cluster of cells was largely replaced by an in-phase cluster. This indicated that only a fraction of cells initiated distinct STAT1/2 dynamics triggered by IFNα from that by IFNβ when exposed to different repeated IFN-I pulses.

Refractory states of signaling proteins have been reported to be involved in inhibition of cellular responses (Vizan *et al*., 2013; Sarasin-Filipowicz et al., 2009). Double-pulse stimulation with different time intervals demonstrated a refractory state of STAT1/2 activity and a desensitization effect on IFNα or IFNβ restimulation, with a limited recovering rate of STAT1/2 activation over time interval. The refractory state may not be stochastic due to its conservation in individual cells for more than 8 hours (Adamson *et al*., 2016). Mathematical modelling predicted that the TNFα refractoriness in NF-κB system was due to pseudo-stable cellular states or levels of signaling proteins in the pathway, as well as associated with negative feedback regulation (Adamson *et al*., 2016). Surprisingly, cells retreated with IFNα showed a biphasic recovering activation rate with increasing time interval, while IFNβ-restimulated cells displayed a monotonically increasing activation rate. These results suggest that upon repeated IFNα exposure, a delayed inhibition on STAT1/2 reactivation occurred. This was suspected to be caused by USP-18, an important negative regulator playing a role in signaling mediated by IFNα but not by IFNβ (Wilmes *et al*., 2015; Arimoto *et al*., 2018; Vizan *et al*., 2013). Therefore, cells displayed distinguished STAT1/2 refractory states upon a double-pulse IFNα input from those by repeated IFNβ stimulation.

In the NF-κB system, cells refractory to TNFα responded to interleukin 1β (IL-1β) (Adamson *et al*., 2016), although it remained unclear whether this would also occur vice versa. This may allow cells to use the refractory states of the NF-κB pathway to initiate robust discrimination between different temporal inputs. Similarly, we suggest that the refractory states in STAT1/2 system might also be able to discriminate between IFNα and IFNβ upon alternate pulses. Although both IFNα and IFNβ evoked STAT1/2 refractory states, stimulation with αβ pulses revealed that cells refractory to IFNα were still able to initiate a strong response to IFNβ, but not vice versa. This could be attributed to higher potency of IFNβ than IFNα, and USP18-based negative feedback control that negatively regulates STAT1/2 signaling by IFNα but not by IFNβ (Wilmes *et al*., 2015; Arimoto *et al*., 2018; Vizan *et al*., 2013). To prevent dysfunction caused by overresponses to cytokines, cells may need to switch to refractory states so as to attenuate their responses. In contrast, when encountering severe infection, cooperation may occur between multiple cell types, leading to alternating waves of different IFN-Is that can readjust the extent of responses in order to make appropriate cell fate decisions.

Overall, we believe that the STAT1/2 system can discriminate between IFNα and IFNβ at diverse temporal conditions, and lead to distinct downstream cell fates. Expanding our knowledge of dynamic STAT1/2 responses to include temporal activation patterns and single-cell heterogeneity are instrumental in developing a comprehensive map of distinct cell-fate decision making in IFN-I signaling and provide an important tool to identify the discriminatory capability of cells in signaling by different ligands.

## Materials and Methods

### Cell line and culture

NIH3T3 cell line stably expressing STAT1-CFP and STAT2-YFP fusion proteins was provided by Mario Köster (Helmholtz Centre for Infection Research, Braunschweig, Germany). The reporter cells were cultured in Dulbecco’s modified Eagle’s medium (DMEM) (Gibco™) supplemented with 10% v/v fetal bovine serum (Gibco™), 1% v/v penicillin-streptomycin (Gibco™), and 2.5 μg/mL puromycin (Gibco™) at 37°C in a humidified 5% CO_2_ incubator. All the above chemicals were purchased from Life Technologies Europe B.V., Bleiswijk, Netherlands.

### Microfluidic device design and fabrication

We employed a previously established microfluidic cell culture platform (Figure 1B) (Yang *et al*., 2022), with a simplified microfluidic device (Figure 1C), for cellular signaling study. The microfluidic device is made of two-layer polydimethylsiloxane (PDMS) with a bottom flow layer for sample loading and a top control layer for valve actuation (Figures 1C and S6). The flow layer comprises a channel network with 4 parallel cell chambers. In the top control layer, we designed a valve array to form a binary tree multiplexer that can be programmed to address each cell chamber (2000 × 500 × 45 μm) individually (Figure S6A). A defined input such as a continuous, single- or double-pulse signal (Figure 1F) can be delivered to specific cell chambers using a custom written MATLAB GUI software (Mathworks, Austin, USA). The experimental workflow includes (Figure 1B): (1) loading cells in multiple chambers, (2) culturing cells until they are stretched, (3) stimulating cells by perfusing medium containing IFN-I (input), (4) monitoring cellular response via time course live-cell imaging, (5) image analysis using CellProfiler pipeline and Python script.

The microfluidic device was fabricated using multilayer soft lithography (Figure S6B) (Yang *et al*., 2022). Briefly, high-resolution film photomasks were printed for the flow layer (microchannels and cell chambers) and the control layer (pneumatic microvalves), separately. Patterns were designed using AutoCAD 2021 (Autodesk GmbH, Munich, Germany), appended as Figure S6A. For the flow layer, a two-level master was prepared on a silicon wafer using a AZ 40XT (MicroChemicals GmbH, Ulm, Germany) and SU-8 3025 photoresist (micro resist technology GmbH, Berlin, Germany) with the aid of a MJB4 mask aligner (SUSS MicroTec Netherlands B.V., Eindhoven, Netherlands). For the control layer, a one-level master was prepared using SU-8 3025 photoresist. Then the two masters were exposed to trichloro(1H,1H,2H,2H-perfluorooctyl)silane (Sigma-Aldrich Chemie N.V., Zwijndrecht, Netherlands) for 2 h. Sylgard 184 PDMS base and curing agent (Dow Chemical Company, Midland, USA) were mixed at a ratio of 5:1 wt/wt, degassed, and decanted onto the master of control layer, and partially cured in a 80°C oven for 20 min. In the meantime, another layer of PDMS mixture with 20:1 PDMS/curing agent ratio was spincoated on the master of flow layer, and incubated at 80°C for 40 min. Two layers of partially cured PDMS were aligned and permanently bonded by heat. After curing, inlet and outlet ports were punched through the structured PDMS layer, with subsequent bonding to a clean glass slide by plasma treatment and incubation in an oven at 80°C for at least 2 h.

### Microfluidic device setup

The control layer of PDMS device was connected to solenoid valves (Festo BV, Delft, Netherlands) with DI-water filled PTFE tubings (Elveflow, Paris, France). We employed the solenoid valves to control pneumatically actuated microvalves on the device including the multiplexer and the microvalves for inlet and outlet. The solenoid valves were managed through the inhouse Matlab script. We closed all microvalves for 10 min to force DI water to fill the control layer. The inlets of flow layer were connected to a pressure pump (Fluigent Deutschland GmbH, Jena, Germany) with sample or chemical filled tubings. The outlet was connected to a waste container. We applied a pressure of 50 kPa to fully perfuse the cell chambers with 100 ng/mL fibronectin solution (Sigma-Aldrich Chemie N.V., Zwijndrecht, Netherlands). In order to de-bubble the device, the microvalve for outlet was closed and air was forced out through gas-permeable PDMS. After 2 hours of incubation, medium was flushed to wash away excess fibronectin in the device.

### Microfluidic cellular signaling experiments

The cultured reporter cells, with 80-90% confluency, were detached from a culture flask with Trypsin-EDTA (Gibco™, Life Technologies Europe B.V., Bleiswijk, Netherlands). The harvested cells were resuspended in 0.5–1.0 mL of fresh DMEM, then loaded in a PTFE tubing, and connected to the microfluidic device. Prior to seeding, the cell sample was filtered using Pre-Separation Filters (30 μm) (Miltenyi Biotec B.V., Leiden, Netherlands) to remove cell clumps. Subsequently, the cells with a concentration of 2×10^6^/mL were injected into each cell chamber through the pressurized tubing controlled by the pressure pump. Once arrived in a chamber, cells were confined by immediate switching off the valves corresponding to the respective chamber. Afterwards, the device was placed at 37°C in a microscope incubator (Okolab S. R. L., Pozzuoli, Italy) with humidified 70% CO_2_. After 3 hours of cultivation, the stretched cells were stained with BioTracker 650 red nuclear dye (Sigma-Aldrich Chemie N.V., Zwijndrecht, Netherlands), followed by stimulation with recombinant mouse IFNα or IFNβ1 (BioLegend Uk Limited, London, UK) at each condition.

### Image data acquisition and analysis

Time-lapse images with a time interval of 7 min were acquired via each fluorescence channel (CFP, YFP and Cy5) using a Nikon Eclipse Ti2 Inverted Microscope equipped with a DS-Qi2 camera (Nikon Instruments Europe B.V., Amsterdam, Netherlands). The XYZ stage served as the microfluidic device holder. Stage motion and image acquisition was controlled via Nikon’s NIS-Elements AR software. We used a 10× objective for setting up the microfluidic device, and a 20× objective for time-lapse image acquisition during experiments.

The acquired image datasets were processed using the in-house CellProfiler pipline. In order to quantify STAT1/2 nuclear translocation dynamics, we extracted the values of nuclear and cytoplasmic fluorescence intensities from tracked individual cells per frame, and subsequently plotted graphs using our created python script.

### Quantitative descriptors of STAT1/2 nuclear translocation dynamics

The duration of a STAT1/2 response peak was defined as a time range starting from the timepoint when an IFN input is applied and ending at the timepoint when the normalized nuclear STAT intensity returns to 1/3 of peak value (1/3 F_max_) (after crossing the peak F_max_) (Figure S8A) (Ryu *et al*., 2015), since STAT trajectories virtually never completely return to the base level. If cells show a sustained activity pattern and do not return to 1/3 of their peak value F_max_, the 1/3 F_max_ will be replaced by F_max_ for calculating duration.

Rate_in_ (slope), the maximal rate of STAT nuclear translocation, was calculated from the dynamic curve as the slope of the straight line fit to three points on the single-cell time course centered at 1/2 of the timepoint when the normalized nuclear STAT intensity reaches maximum (Figure S8B) (Zhang *et al*., 2017).

AUC (Area under the curve) was calculated by measuring the area bounded by the nuclear STAT time course and the baseline amount of nuclear STAT (Figure S8C). Negative values were regarded as zero and therefore was discarded from analysis.

### Determination of STAT1/2 activation threshold

To define the thresholds for determining STAT activation status, we calculated an average fold change value for nuclear STAT1 and STAT2, respectively, from each single cell in a limited number of time course images acquired from blank control experiments, where IFN-I was replaced by PBS. We further calculated the fold change threshold value based on a 95% tolerance interval (Figure S9A and B), which was defined as STAT1/2 activation threshold. In continuous or single-pulse experiments, a cell showing an average STAT fold change value (based on the first few frames) above this threshold value was determined as active, otherwise as negative.

Given that cells may still be active when subjected to a second-pulse stimulation, a corrected STAT fold change value of each cell from blank control data was calculated by F’_max_/F’_i_, where F’_max_ is the peak intensity in response to the second-pulse stimulation, and F’_i_ is the STAT intensity when the second-pulse IFN-I is applied. Similarly, the threshold value for determining STAT activation after a second pulse stimulation was calculated based on a 95% tolerance interval (Figure S9C).

### Clustering of STAT1/2 dynamics

The algorithms we used for clustering STAT dynamics were all supervised clustering algorithms, i.e., we defined the number of clusters ourselves, since unsupervised clustering algorithms could result in over or under segmentation of single-cell trajectories (Aghabozorgi et al., 2015). We used five different clustering algorithms to evaluate their performance in our data analysis, including K-means clustering with three different methods of calculating distances (Euclidean, dynamic time warping (DTW) and soft-DTW), self-organizing map (SOM) (Kohonen, 1998), and Hierarchical clustering algorithms (Figure S10A and B). SOM was selected as the optimal clustering algorithm based on clustered single-cell trajectories displayed in heatmaps and Pearson correlation coefficients, and was thus applied to cluster STAT dynamics (Figure S10A and C).

## Acknowledgements

We are grateful to Ulfert Rand, Hansjörg Hauser and Mario Köster for providing the STAT1-CFP/STAT2-YFP dual reporter NIH3T3 cell line. This result is part of a project, ImmunoCode, that has received funding from the European Research Council (ERC) under the European Union’s Horizon 2020 research and innovation programme (Grant agreement No. 802791). Furthermore, we acknowledge generous support by the Eindhoven University of Technology.

## Author contributions

**Haowen Yang:** Conceptualization, Methodology, Investigation, Data curation, Formal Analysis, Visualization, Validation, Project administration, Supervision, Writing – original draft, Writing – review & editing; **Thomas van de Kreek:** Investigation, Software, Data curation, Formal Analysis, Visualization, Validation; **Laura C. Van Eyndhoven:** Resources, Writing – review & editing, **Jurjen Tel:** Conceptualization, Resources, Supervision, Funding Acquisition, Writing – review & editing.

