## Supplemental Information for "Temporal perturbation of STAT1/2 activity reveals dynamic ligand discrimination of type I interferon signaling"

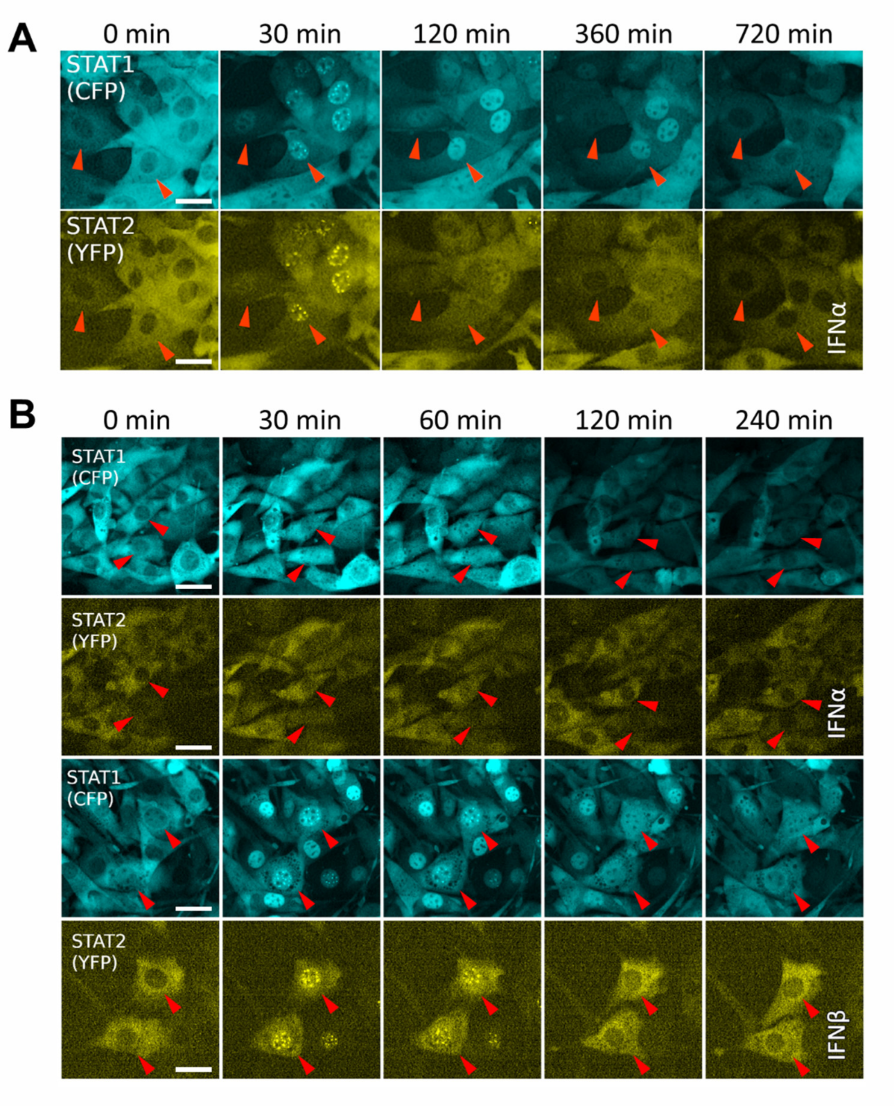


**Figure S1.** Time-lapse images of STAT1-CFP and STAT2-YFP fusion proteins stably expressed in NIH3T3 fibroblasts exposed to (A) a continuous/sustained input of 100 ng/mL IFNα, (B) a single 1 min pulse of 100 ng/mL IFNα or IFNβ.

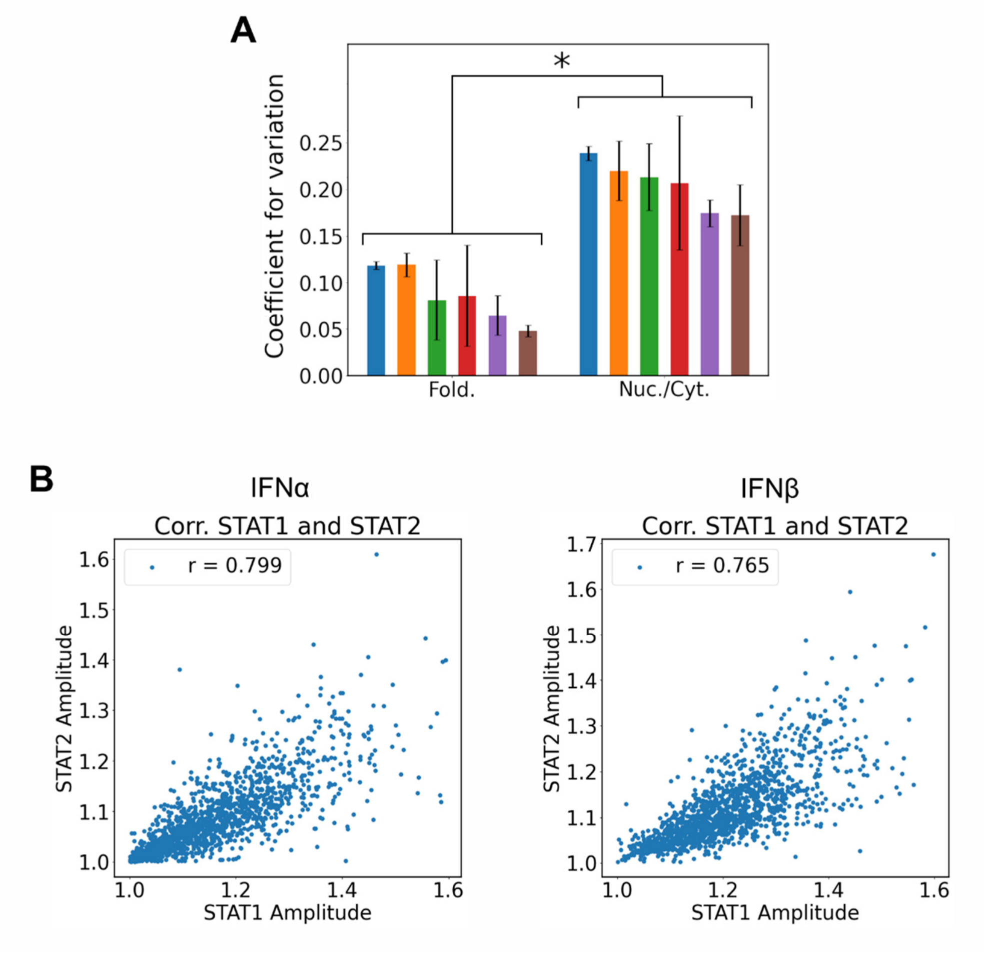


**Figure S2.** (A) Bar graphs of the coefficient of variation for fold change (of nuclear STAT1 intensity) and nuclear/cytoplasmic ratio (of STAT1 intensity). (B) Correlations between STAT1 and STAT2 amplitudes upon IFNα or IFNβ stimulation.

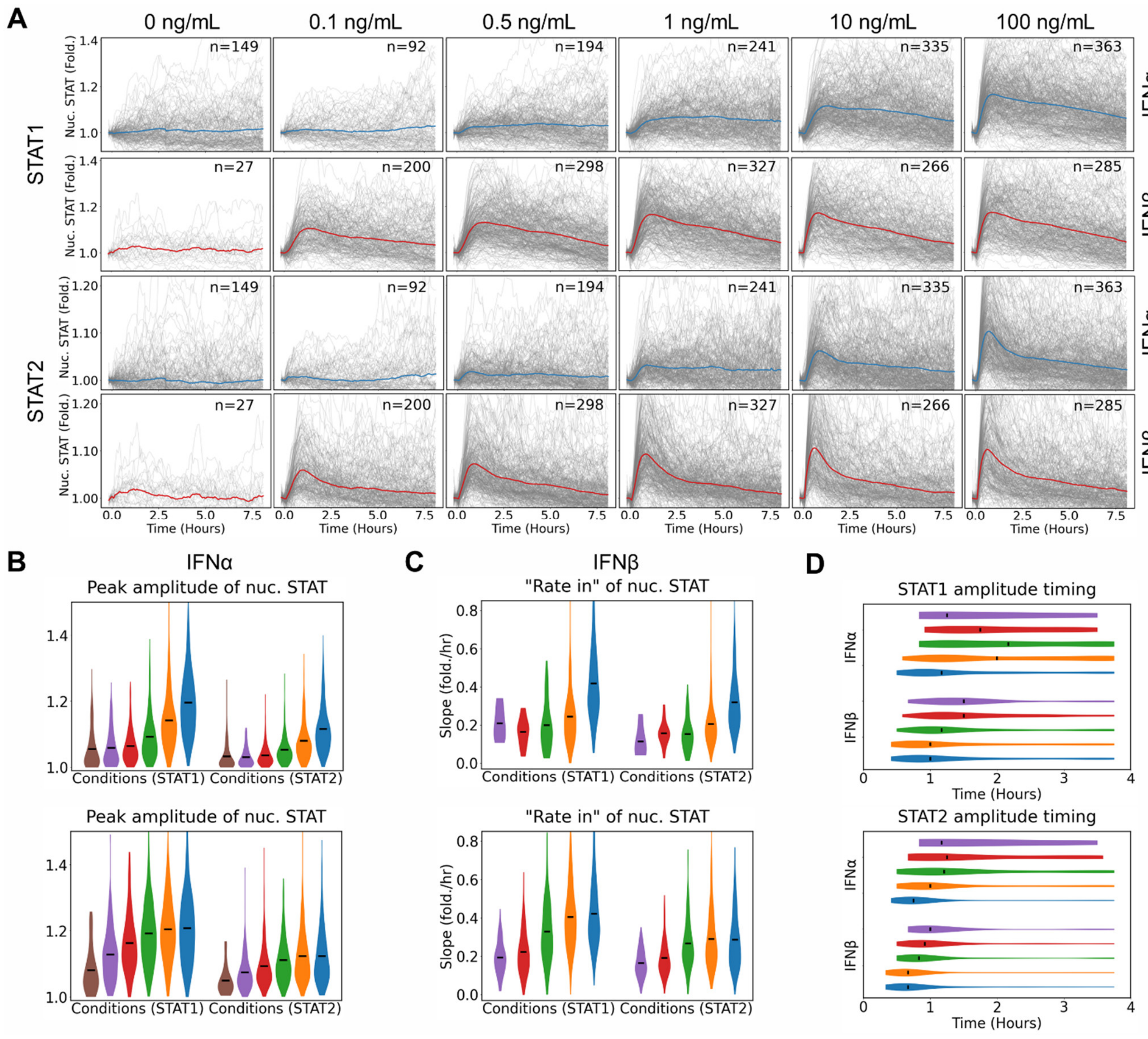


**Figure S3.** (A) Single-cell STAT1 and STAT2 nuclear translocation dynamics upon continuous stimulation with different concentrations of IFNα and IFNβ ranging from 0.1–100 ng/mL (0 ng/mL as a blank control), respectively. STAT1/2 peak amplitudes (B), $Rate_{in}$ (C), and peak amplitude timing (D) of single-cell STAT1/2 trajectories under the conditions in (A).

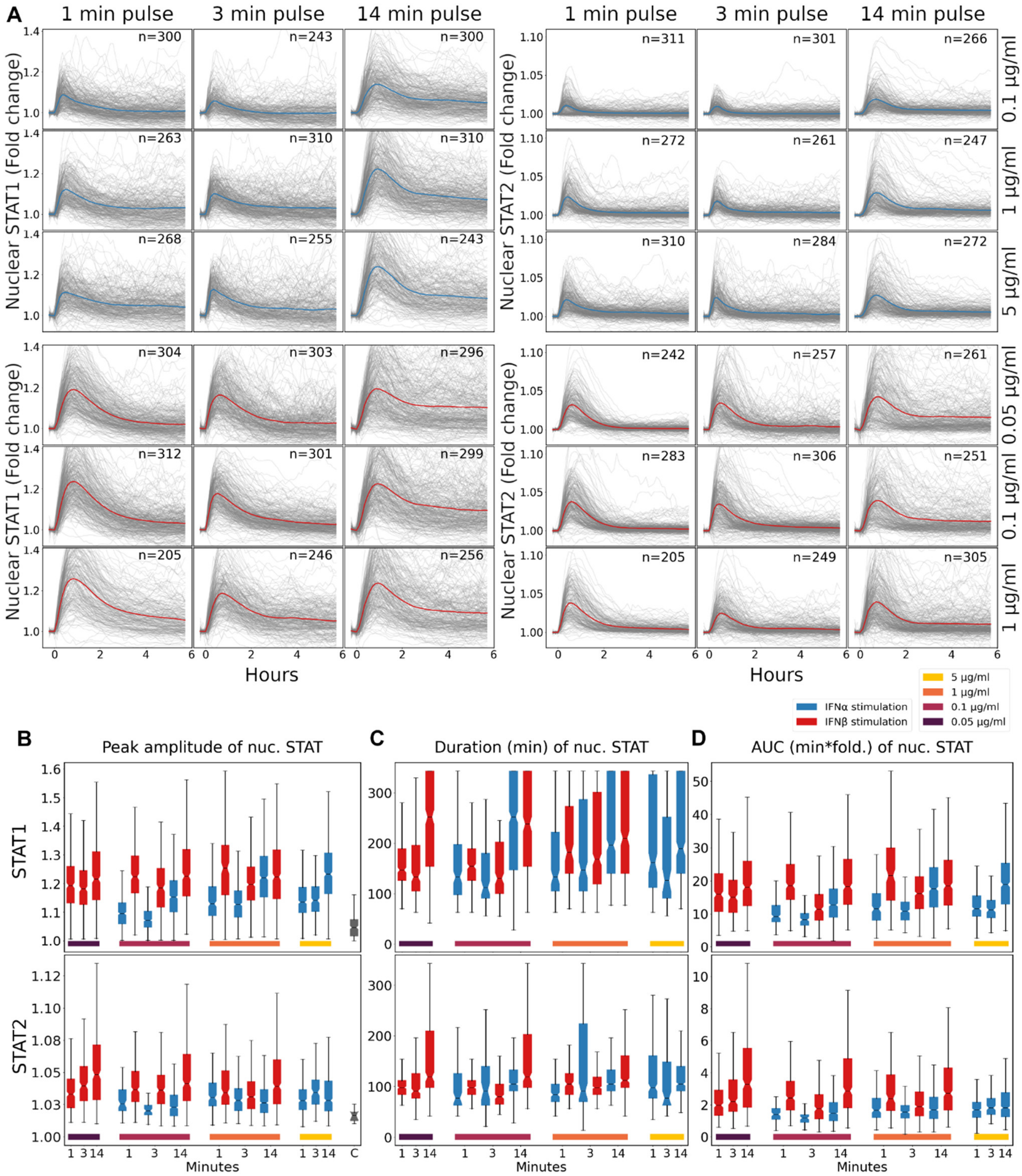


**Figure S4.** (A) Single-cell STAT1 and STAT2 activity trajectories upon single-pulse IFNα/IFNβ stimulation at different concentrations (0.1, 1 and 5 μg/mL) or durations (1, 3, and 14 min). STAT1/2 peak amplitudes (B), Duration (C), and AUC (D) of single-cell STAT1/2 trajectories under the conditions in (A).

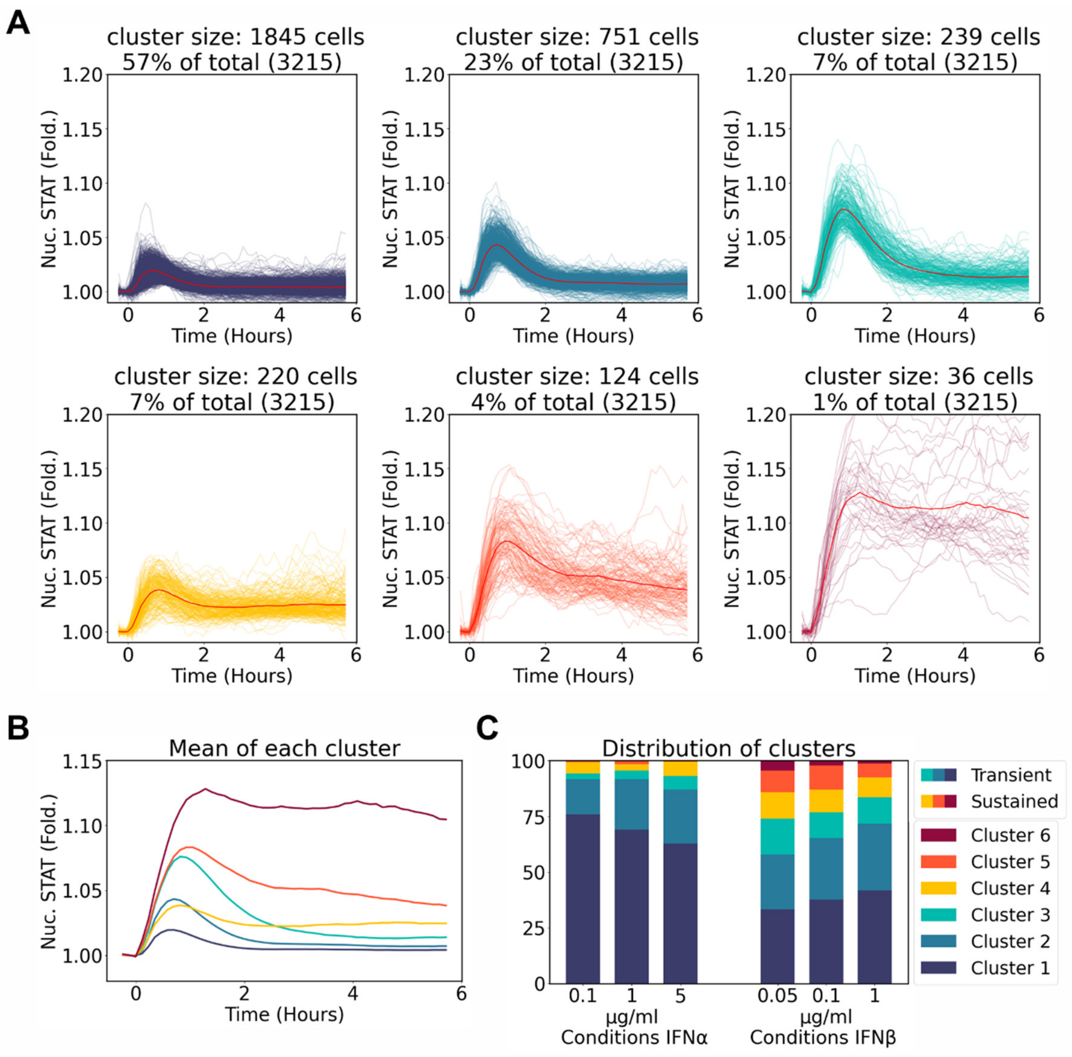


**Figure S5.** (A) STAT1/2 activity trajectories from all cells stimulated with a 14 min IFNα/IFNβ pulse were pooled and 6 clusters were identified. (B) Average STAT1/2 activity across 6 clusters identified from (A). (C) Distribution of STAT1/2 activity trajectories in response to different IFN-I pulse intensities.

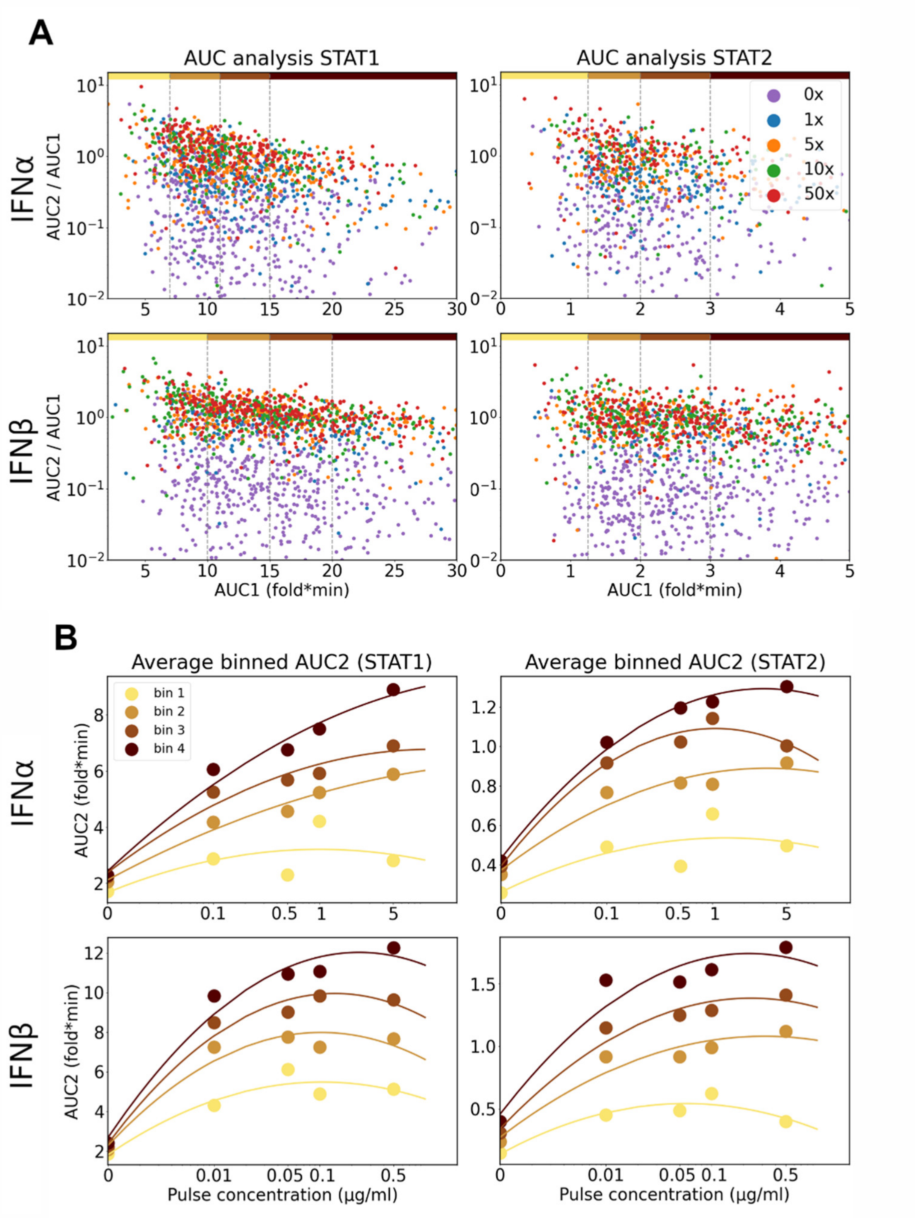


**Figure S6.** (A) Scatterplots of AUC2/AUC1 stratified along AUC1 across the range of pulse conditions. Colored bar along top illustrates bins of single cells based on AUC1 into an approximately equal number of cells per condition. (B) Average responses of single cells (AUC2) to increasing pulse concentrations (0.01, 0.05, 0.1, or 0.5 μg/mL) for each bin in (A).

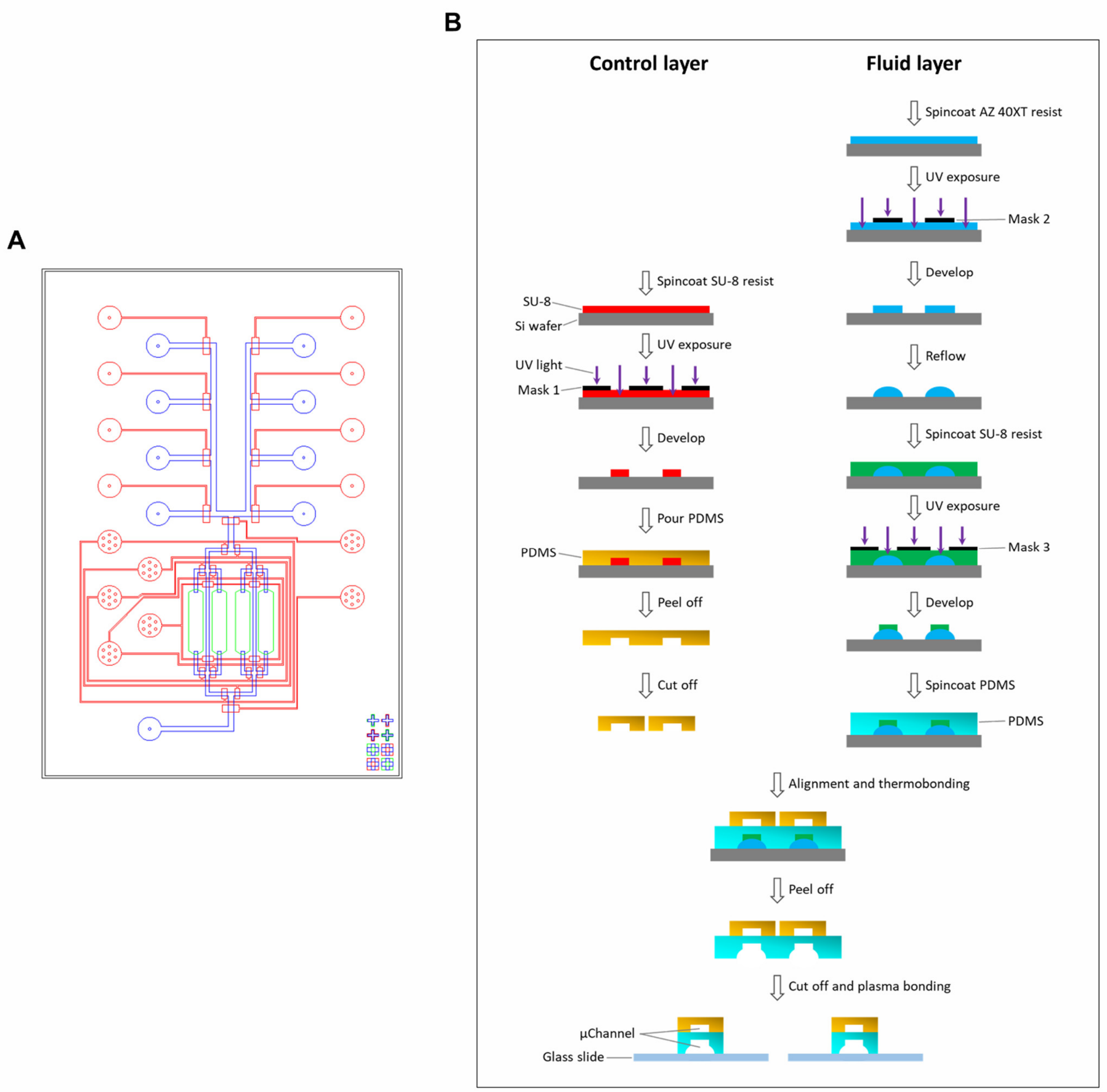


**Figure S7.** (A) Multiple-layer microfluidic device design. The 8 valves at top and the binary tree multiplexer in the middle are pneumatically actuated by the control layer (red). The cells and chemicals are delivered through the flow channels (blue). The parallel 4 chambers (green) where cells are cultured and stimulated are connected between fluid channels. (B) Schematic of the device fabrication routine. For the control layer, SU-8 3025 photoresist was spincoated on a silicon wafer and then patterned through a film photomask. After a post-exposure bake, the SU-8 structures were developed and stabilized with a hard bake. For the two-level fluid layer, AZ 40XT photoresist was used and patterned, developed on a silicon wafer, followed by overnight reflow to make AZ structures round. Afterwards, SU-8 3025 resist was spincoated on this AZ patterned wafer. The second-level structures were generated after a post-exposure bake and development. Upon silanization on both masters, PDMS was poured on the master of control layer, while the master of flow layer was spincoated with PDMS. When partially cured, PDMS is peeled off from the master for control layer and cut off into small pieces. The PDMS of control layer was aligned and thermally bonded with the structures on the flow layer. Once completely cured, the PDMS with entire structures was permanently bonded with a glass slide using plasma bonding.

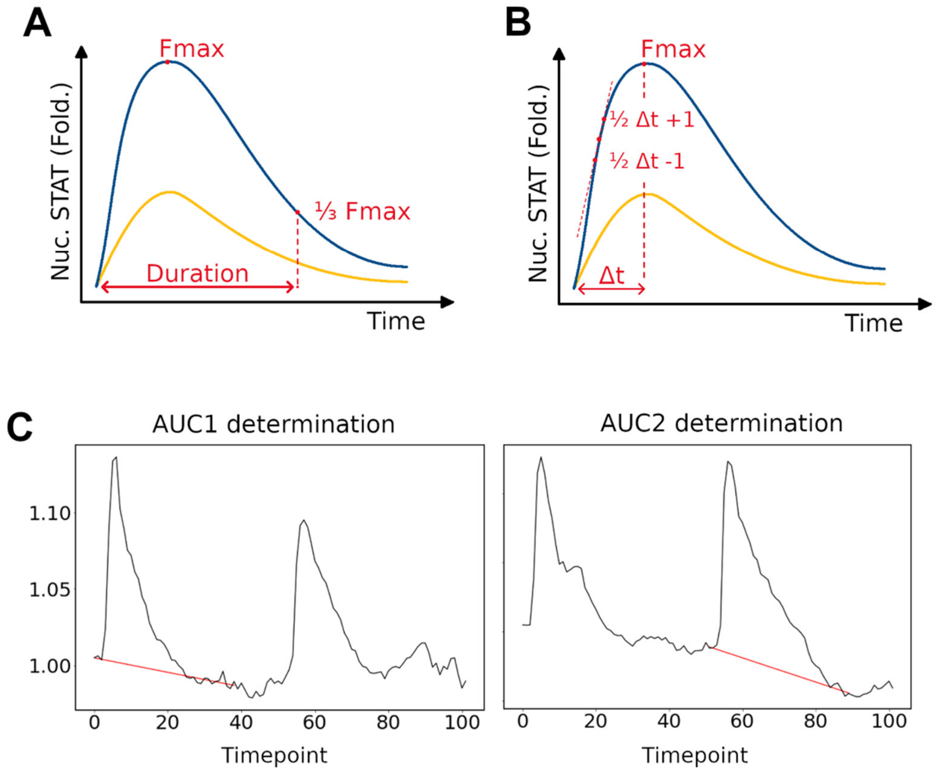


**Figure S8.** Quantitative descriptors of STAT activity trajectory. (A) Duration. (B) $Rate_{in}$. (C) AUC.

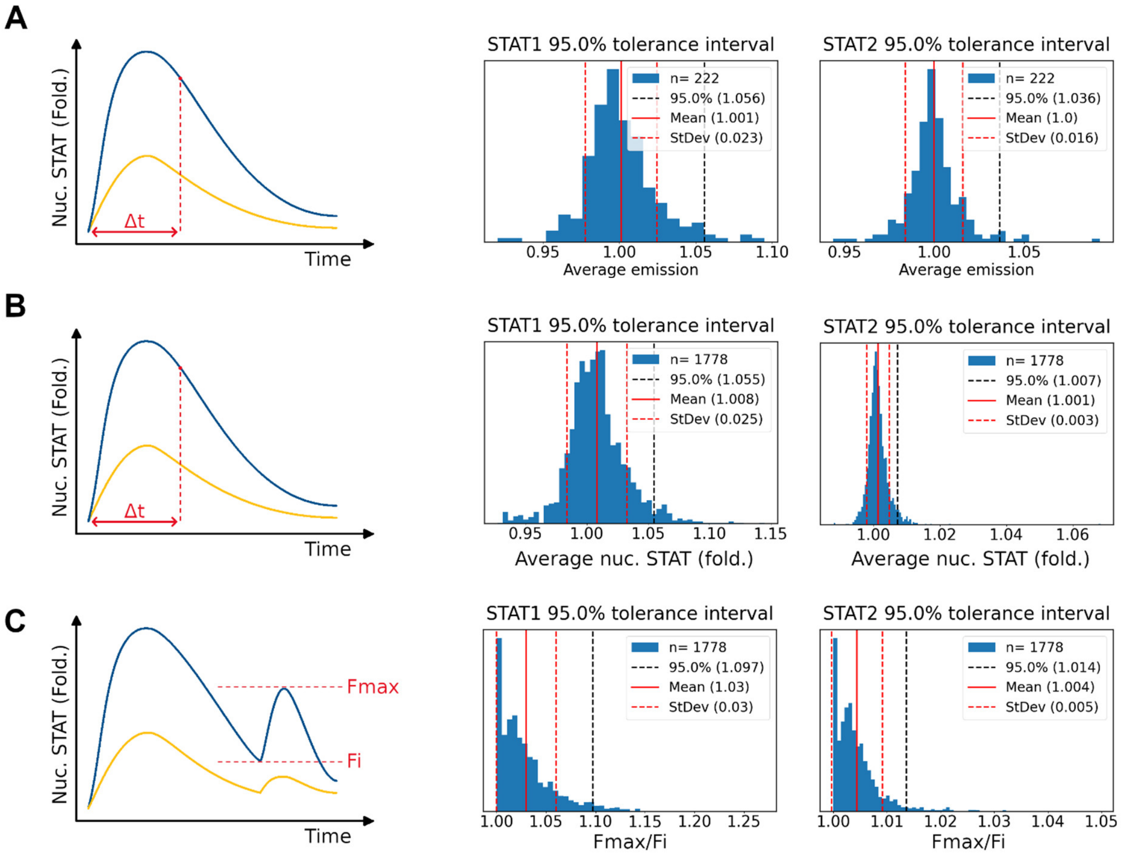


**Figure S9.** Determination of STAT activation threshold in (A) activation upon continuous/sustained stimulation, (B) activation upon single-pulse stimulation, and (C) reactivation after second-pulse stimulation. The average STAT fold change value of each cell is calculated based on the first few frames (Δt).

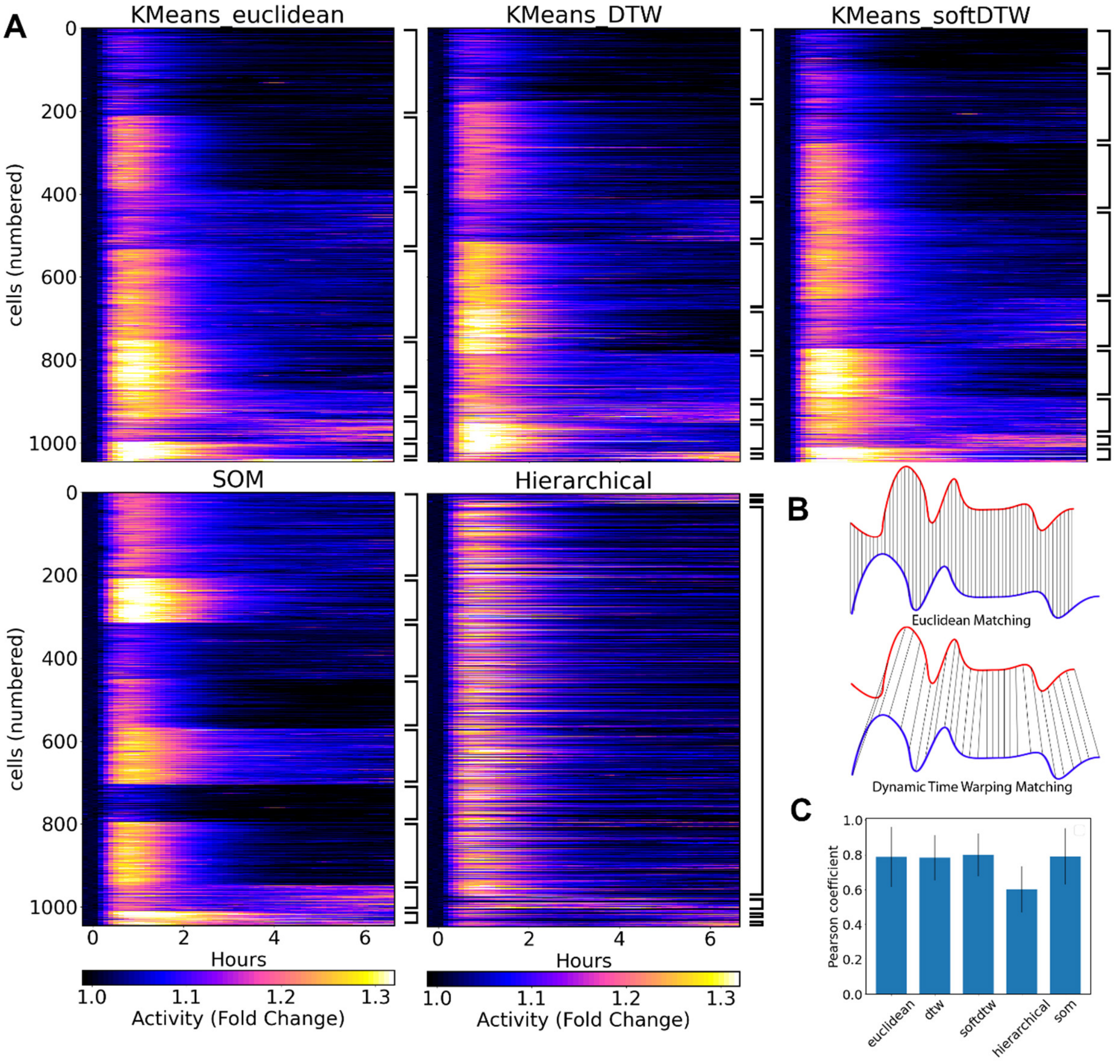


**Figure S10.** (A) Clustered single-cell STAT activity trajectories displayed in heatmaps using different clustering algorithms, including K-means clustering with 3 different methods for distance calculation (Euclidean, dynamic time warping (DTW) and soft-DTW), SOM, and Hierarchical clustering algorithms, respectively. (B) Methods (Euclidean and DTW) for calculating distances between single-cell STAT trajectories. (C) Pearson correlation coefficients between clusters using different clustering algorithms.
